# Resident T cells orchestrate adipose tissue remodeling in a site peripheral to infection

**DOI:** 10.1101/2022.02.24.481701

**Authors:** Agnieszka M. Kabat, Alexandra Hackl, David E. Sanin, Patrice Zeis, Katarzyna M. Grzes, Francesc Baixauli, Ryan Kyle, George Caputa, Joy Edwards-Hicks, Matteo Villa, Nisha Rana, Jonathan D. Curtis, Angela Castoldi, Jovana Cupovic, Leentje Dreesen, Maria Sibilia, J. Andrew Pospisilik, Joseph F. Urban, Dominic Grün, Erika L. Pearce, Edward J. Pearce

## Abstract

Infection with helminth parasites can affect adiposity, but underlying mechanisms that regulate this process are unclear. We found that fat content of mesenteric adipose tissue (mAT) declined in mice during infection with gut-restricted parasitic worms. This was associated with the accumulation of metabolically activated, immunostimulatory cytokine- and extracellular matrix-secreting multipotent stromal cells, which had potential to differentiate into preadipocytes. Concomitantly, mAT became infiltrated with Th2 lymphocytes that took up long-term residence and responded to signals from stromal cells by producing stromal cell-stimulating cytokines, including Amphiregulin. Signals delivered by Amphiregulin to stromal cells were required for immunity to infection. Our findings reveal intricate intercellular communication between Th2 cells and adipocyte progenitors and link immunity to intestinal infection to T cell-dependent effects on the adipocyte lineage.

---

Type 2 immune processes underlie aspects of tissue homeostasis. This is particularly well recognized in adipose tissue (AT), where eosinophils and type 2 innate lymphoid cells (ILC2) contribute to the maintenance of alternative macrophage activation, and are implicated in AT browning and the prevention of metabolic disease (*1–4*). Regulatory T (Treg) cells are also important for AT homeostasis (*5, 6*). Recently, stromal cells (SC) within visceral white AT were shown to produce IL-33, which sustains ILC2, eosinophil and Treg populations during homeostatic conditions (*7–10*). Infection with parasitic helminths drives type 2 immune responses, and *H. polygyrus* infection has been reported to prevent obesity in mice fed a high fat diet, through an IL-33-dependent pathway linked to AT browning (*11, 12*). Visceral AT was shown to undergo functional repurposing towards the production of antimicrobial molecules during inflammatory type 1 immune responses associated with disseminated infection with foodborne microbial pathogens (*13*). However, a detailed understanding of how AT changes during helminth infections is lacking. Recent work using single cell RNA sequencing allowed the identification of a lineage hierarchy of adipocyte progenitors, anchored in a population of MPC marked by *Dpp4* expression (*14*). These MPC are depleted during insulin resistance and obesity, signaling their importance for maintaining AT homeostasis (*14*). We reasoned that immune responses to enteric parasites may affect lineage commitment within the mAT SC populations as a part of the tissue response to inflammation, and that this might account for changes in adiposity associated with helminth infection.

## Results

### Dynamic changes occur in adipose tissue peripheral to the site of infection

*H. polygyrus* infection is chronic, but can be cleared by anthelminthic treatment, after which mice show increased resistance to secondary challenge (Fig. S1). We found that infection-induced mesenteric lymph node (mLN) enlargement was accompanied by a reduction in surrounding mAT (Fig. 1A), characterized by a decrease in mAT weight and fat content (Fig. 1B, C). This coincided with an increase in cellularity of the stromal vascular fraction (SVF), particularly after secondary infection, due to increases in immune and stromal cells in this compartment (Fig. 1D-F).

**Fig 1.**
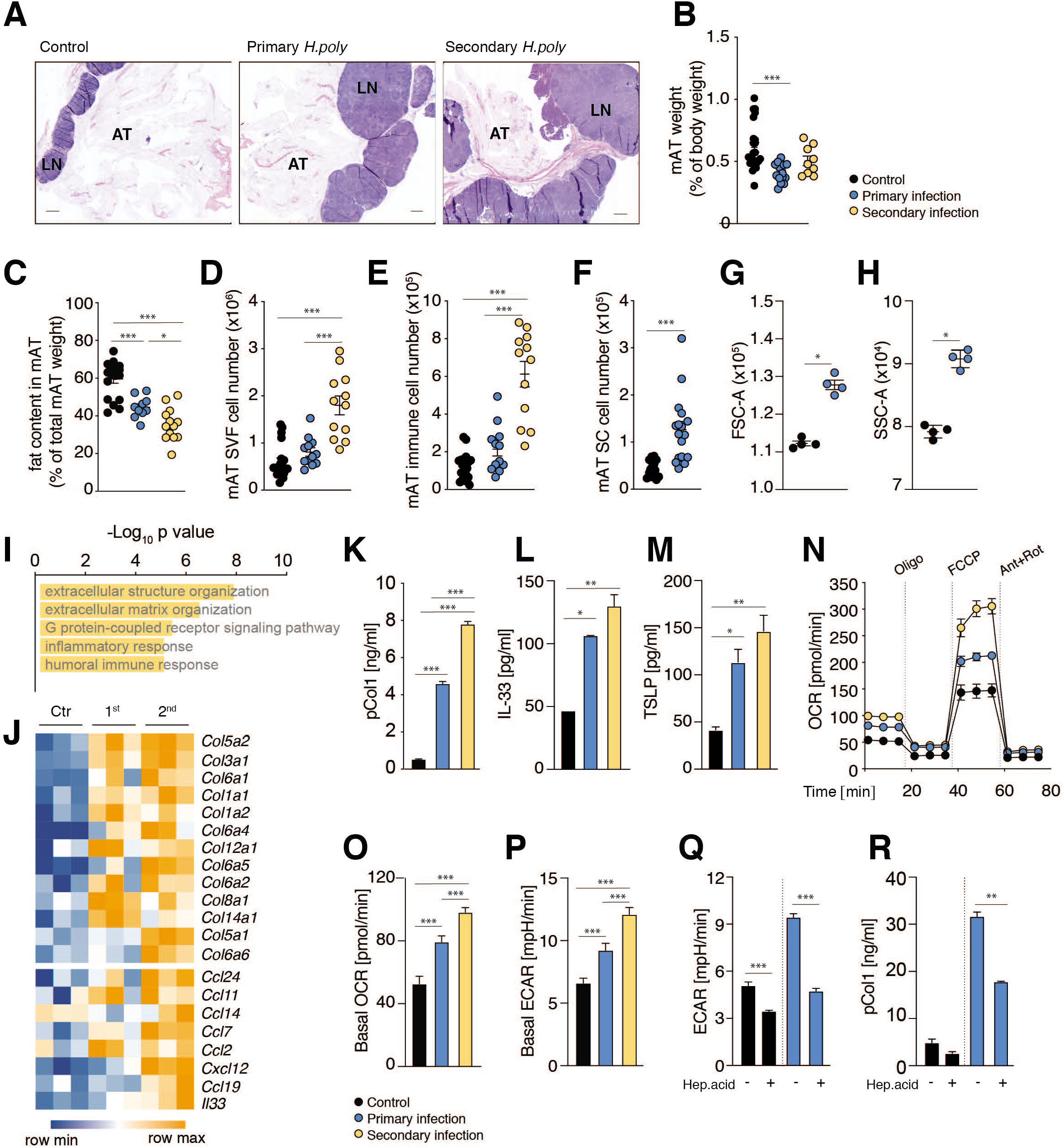
mAT SC ECM production and metabolic rates are upregulated during *H. polygyrus* infection. **(A)**. H&E staining of mAT and mLN during *H. polygyrus* infection, scale bar 500μm (LN-lymph nodes, AT – adipose tissue). (**B**) Weight of mAT of normalized to total body weight. (**C**) Fat content in mAT measured by MRI and normalized to the total mAT weight. (**D-F)**. Total SVF cell counts (D), immune cell counts (gated on live CD45^+^ CD31^-^ cells) (E) and SC counts (gated on live CD45^-^ CD31^-^ PDGFR*α*^+^ Sca1^+^) (F) (**G, H**) Cell size (G) and granularity (H) of SC from mAT of control and infected mice. (**I**) GO enrichment analysis of significantly upregulated genes in SC from control vs primary infections. (**J**) Heatmap showing collagen, cytokine and chemokine expression pattern in SC isolated from mAT of control, and infected mice. (**K-M**) Pro-collagen 1 (pCol1) (K), IL-33 (L) and TSLP (M) production by mAT SC of control and infected mice. (**N**) Oxygen consumption rates, OCR, of mAT SC from indicated conditions at baseline and after Oligomycin (Oligo), FCCP and Rotenone/Antimycin (Rot/Ant) injections. (**O**) Baseline OCR in mAT SC. (**P**) Baseline extracellular acidification rates, ECAR, of mAT SC. (**Q, R**) Effects of heptelidic acid [10μM] on baseline ECAR (Q) and pCol1 production (R) of mAT SC of control and infected mice. Dots represent biological replicates (BR), error bars represent SEM from BR (B-H) or technical replicates (TR) (K-R). Data combined from 2-3 independent experiments (B-F), representative of 2-3 experiments (A, G, H, K-R), or from one experiment (I, J).

We noted that infection caused an increase in stromal cell size and side scatter, indicating that the cells had become activated and potentially secretory (Fig. 1G, H). To understand infection-associated changes in mAT, we performed RNAseq on sorted CD45^-^ CD31^-^ Sca1^+^ PDGFR*α*^+^ SC. Pathway enrichment analysis revealed an emphasis on ECM and inflammation with expression of collagen, chemokine and cytokine genes during infection (Fig. 1I, J). Consistent with this, mAT SC isolated from infected mice constitutively secreted pro-collagen 1 (pCol1), TSLP and IL-33 when cultured *ex vivo* (Fig. 1K-M).

The increased size, granularity and secretory activity of SC from infected mice suggested an increase in anabolic metabolism. This was confirmed by *ex-vivo* extracellular flux analysis, where mAT SC from infected mice had increased oxygen consumption rates (OCR) indicative of increased mitochondrial respiration (Fig. 1N, O) and increased extracellular acidification rates (ECAR) (Fig. 1P), indicative of increased production of lactate from glycolysis. OCR and ECAR paralleled the secretion of collagen and cytokines, being more pronounced after secondary infection (Fig. 1O, P). Metabolic activation was important for altered stromal cell function since the selective glycolysis inhibitor heptelidic acid, which targets GAPDH, inhibited pCol1 production in parallel with reducing ECAR (Fig. 1Q, R).

### Th2 cells with innate immune cell properties take up long term residence in adipose tissue

We explored infection-induced changes in the cellular composition of the SVF in greater detail using single cell (sc) RNAseq. Unsupervised clustering of SVF cell transcriptomes revealed multiple cell clusters (Fig. 2A, B, Fig. S2A). Striking infection-induced changes within the immune cell compartment were apparent, with a reduction in myeloid cells and an expansion of the CD4^+^ T cell compartment (Fig. 2B-D). These changes were confirmed by flow cytometry (Fig. 2E, F), which also revealed an increase after infection of mAT eosinophils (Fig. 2G), that was not detected by scRNAseq. The majority of mAT CD4^+^ T cells in infected mice were GATA3^+^ and therefore Th2 cells (Fig. 2H, S2). mAT Th2 cells were characterized by high expression of receptors for the stroma derived cytokines IL-33 and TSLP, whereas these cells constituted only a small percentage of mLN cells (Fig. 2H-J, Fig. S3A, B). Indeed, in secondary infection up to 80% of all mAT CD4^+^ T cells were GATA3^+^ IL-33R^+^ Th2 cells (Fig. 2I). We found that mAT Th2 cells were CD69^+^ CD44^+^ and CD62L^-^, suggesting that they were resident memory T cells (T_RM_) (Fig. S3C). Prior studies located AT resident T cells in tertiary lymphoid structures, also called fat associated lymphoid clusters (FALCs) (*13, 15, 16*). In line with this, whole mount mAT confocal microscopy analysis revealed numerous FALCS enriched in CD3^+^ GATA3^+^ (Th2) cells in the infected mice (Fig. S4A, B). These structures were rare and smaller in mAT from uninfected mice. Moreover, in infected mice Th2 cells were also present in areas outside of the FALCs, scattered among Perilipin-1^+^ adipocytes and PDGFRa^+^ SC (Fig. S4B, C).

**Fig 2.**
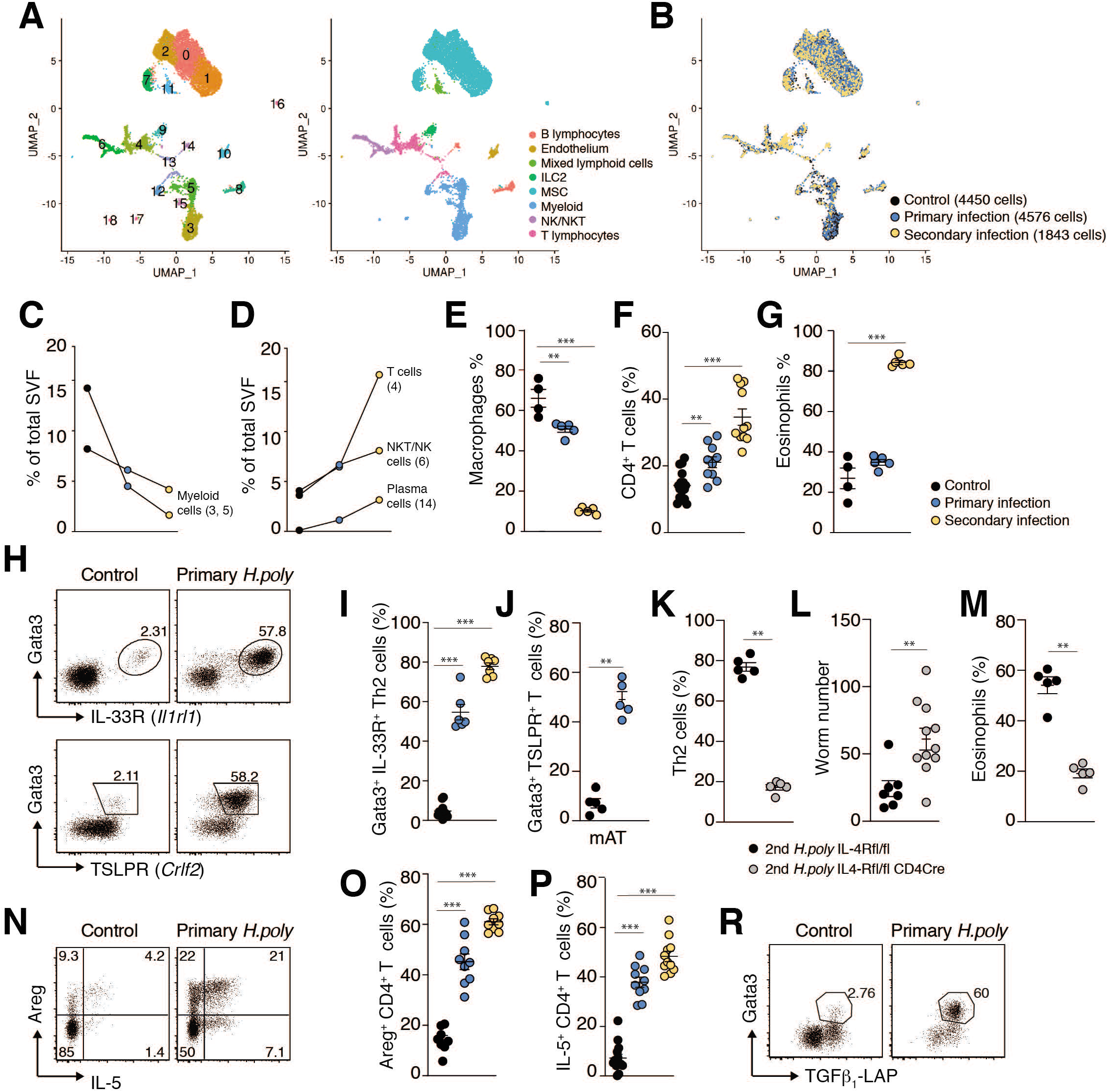
Th2_RM_ cells dominate the mAT T cell population in *H. polygyrus* infection. (**A, B**) Uniform Manifold Approximation and Projection (UMAP) plots of 10869 SVF cells isolated from mAT depots of control, primary infected and secondary infected mice (one mouse per condition). Unsupervised clustering identified 18 cell groups; plots are color-coded according to cell cluster and broad identification of cell types (A) or experimental condition (B). (**C, D**) Changes in immune cell populations after infection (normalized to total cell number in experimental condition). Number in brackets indicate cluster ID from (A). **(E-G**) Frequencies of macrophages (F4/80^hi^ SiglecF^low^ CD11b^+^) (E), CD4^+^ TCRβ^+^ T cells (F) and eosinophils (F4/80^low^ SiglecF^hi^ CD11b^+^) (G) in mAT in control and infected mice. (**H-J**) Representative FACS plots (H) and quantification (I, J) of GATA3^+^ IL-33R^+^ and GATA3^+^ TSLPR^+^ Th2 cells (gated on Foxp3^-^CD4^+^ TCRβ^+^ CD45^+^ live cells in lymphocyte gate) in mAT during infection. (**K-M**) *Il4ra*^fl/fl^ *Cd4*-*Cre* and *Il4ra*^fl/fl^ mice after secondary infection. Frequencies of Th2 cells (IL-33R^+^ GATA3^+^ FOXP3^-^CD4^+^ TCRβ^+^) (K), eosinophils (M) (SiglecF^hi^ SSC-A^hi^ CD45^+^) and worm numbers in small intestine (L). (**N-P**) Representative FACS plots (N) and quantification of Areg (O) and IL-5 (P) expression by mAT CD4^+^ T cells (as proportion of CD4^+^ TCRβ^+^ Foxp3^-^ CD45^+^ live cells in lymphocyte gate) during infection. (**R**) TGFβ_1_-LAP expression by mAT Th2 cells in control and infected mice (gated on Foxp3^-^ CD4^+^ TCRβ^+^ CD45^+^ live cells in lymphocyte gate). Dots represent BR and error bars represent SEM from BR (E-G, I-M, O, P). Data combined from 2-3 independent experiments (F, I, O, P), representative of 2-3 experiments (E, G, J-M), or from one experiment (A-D, R).

Prevention of Th2 cell development through the deletion of IL-4R*α* on T cells in *Il4ra*^fl/fl^ *Cd4*-Cre mice resulted in a reduction of resistance to secondary infection, and the loss of associated components of the response such as tissue eosinophilia (Fig. 2K-M), confirming the importance of Th2 cells for host defense during secondary infection. Functionally, the majority of mAT GATA3^+^ Th2 cells were capable of making the eosinophil survival factor IL-5, but also the tissue modulatory cytokines Areg and TGFβ_1_ (Fig. 2N-R, Fig S5A, B); cells with these attributes were rare in the mLN from the same animals (Fig. S3C, D), consistent with the view that terminal Th2 cell differentiation occurs within peripheral tissues (*17*). scRNAseq data indicated that during infection Th2_RM_ cells became the main producers of Areg and TGFβ_1_ in the mAT, eclipsing the contribution of ILC2 (Fig. S5E). Lastly, we found that the increase in Th2_RM_ population was paralleled by the progressive infection-associated decline in mAT Treg cells and IFN-*γ* producing CD4^+^ T cells (Fig. S5F, G), which was not apparent in infected *Il4ra*^fl/fl^ *Cd4*-Cre mice (Fig. S5H), indicating that Th2 cells compete with Treg and Th1 cells for the available mAT niche.

To ask whether mAT Th2 cells have tissue-specific attributes, we used scRNAseq to compare them to Th2 cells sorted from anatomically related sites affected by infection, namely the small intestine (SI) and mLN. Clustering and similarity analysis with VarID (*18*) revealed distinct groupings of T cell populations by tissue of residence (Fig. 3A,B). *Cd44^+^*, *Cd69^+^*, *Cd62l^-^* (*Sell*), *Klf2^-^* Th2_RM_ cells were present in mAT and SI but nevertheless clustered separately from each other (C7 vs. C11) (Fig. 3A, C), indicating location-dependent functional distinctions. Previous work has described Th2_RM_ cells in SI and the peritoneal cavity of *H. polygyrus* infected mice, but not in mAT (*19*). We observed that mAT, but not SI Th2_RM_ cells, upregulated the expression of CD25 (*Il2ra*), and expressed high levels of *Il1rl1* (IL-33R, ST2), while SI Th2_RM_ cells displayed higher expression of *Il17rb* (IL-25R), reminiscent of tissue-specific alarmin receptor expression in ILC2 (*20*) (Fig. 3D). Furthermore, Th2_RM_ cells from mAT, to a greater extent than SI Th2_RM_ cells, expressed several genes previously associated with ILC2: *Nmur1, Calca (or Cgrp), Klrg1 and Arg1,* indicating that Th2_RM_ cells acquire an “innate-like” phenotype when residing in mAT (Fig. 3D). In contrast to SI Th2_RM_ cells, mAT Th2_RM_ cells also expressed *Ccr2*, pointing towards a possible role for CCR2 ligands in Th2_RM_ cell localization in mAT (Fig. 3E). Moreover, analysis of integrin expression patterns by both scRNAseq and flow cytometry revealed that mAT Th2_RM_ cells expressed integrins that enable them to interact with ECM (Fig. 3E, Fig S6A). Whereas the mAT Th2_RM_ cell cytokine signature was dominated by *Il5, Il6, Areg* and *Tgfβ_1_,* SI cells additionally strongly expressed *Il4* and *Il13* (Fig. 3F).

**Fig 3.**
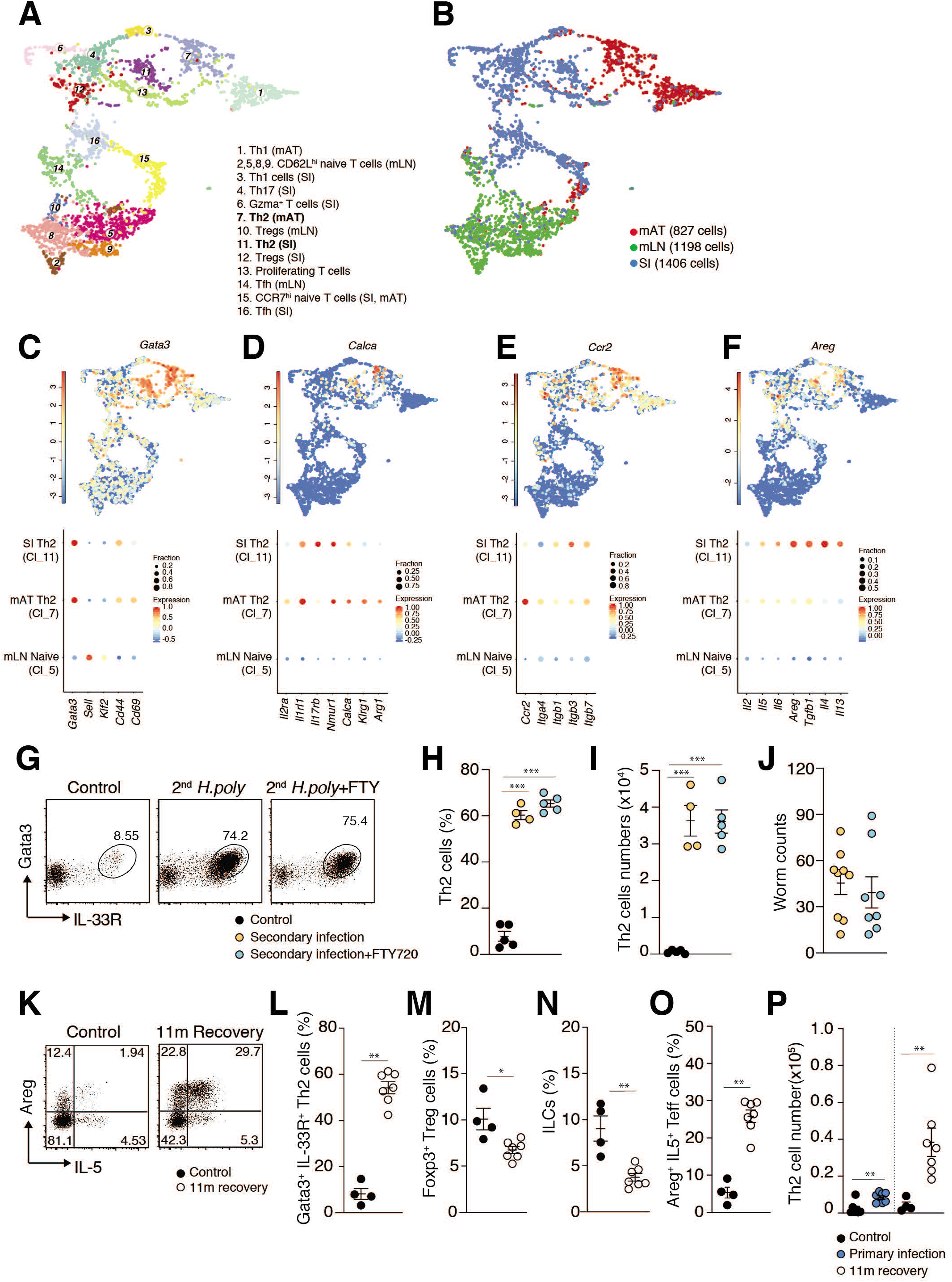
Permanent remodelling of the mAT lymphoid niche after *H. polygyrus* infection. (**A, B**) UMAP plots of CD4^+^ TCRβ^+^ T cells sorted from mLN, mAT and SI of control, primary and secondary infected mice. Unsupervised clustering distinguished 16 cell groups; plots are color-coded according to cell identity (A) or tissue of origin (B). (**C-F**). UMAP plots indicating log2 normalized expression of selected gene (upper panels) and cluster-specific gene expression shown as dot plots (lower panels), where color represents the z-score of the mean expression across clusters and dot size represents the fraction of cells in the cluster expressing the selected gene. (**G-J**). Mice were subjected to secondary infection and treated with FTY720 throughout the infection where indicated. Representative FACS plots (G), percentages (H) and numbers (I) of GATA3^+^ IL-33R^+^ Th2 cells (gated on Foxp3^-^ TCRβ^+^ CD45^+^ live cells in lymphocyte gate). Worm counts from small intestine (J). (**K, O**). Representative FACS plot (K) and frequencies (O) of mAT IL-5^+^ Areg^+^ T cells (gated on Foxp3^-^ CD4^+^ TCRβ^+^ CD45^+^ live cells in lymphocyte gate). (**L-N**) Frequencies of mAT GATA3^+^IL-33R^+^ Th2 cells (gated on ^+^ Foxp3^-^ CD4^+^ TCRβCD45^+^ live cells in lymphocyte gate), Foxp3^+^ Treg cells (gated on CD4^+^ TCRβ^+^ CD45^+^ live cells in lymphocyte gate) and ILC (gated on Lin-CD45^+^ Thy1^+^ live cells in lymphocyte gate; see Methods). (**P**) Comparison of Th2 numbers in mAT during primary infection and 11 months recovery. Dots represent BR and error bars represent SEM from BR (H-J, L-P). Data combined from 2-3 independent experiments (A-F, J, P) or representative of 2-3 experiments (G-I, K-O).

Consistent with mAT Th2 cells being T_RM_, Th2 cell accumulation during secondary infection was unaffected by treatment with FTY720, a sphingosine-1-phosphate receptor 1 agonist (Fig. 3G-J), indicating that the expansion of the mAT Th2_RM_ population is independent on the recruitment of cells from secondary lymphoid organs. Furthermore, FTY720 treatment had no effect on resistance to reinfection (Fig. 3K). It was striking that mAT remained enriched for IL-33R^+^ Th2_RM_ cells capable of making both IL-5 and Areg for up to 11 months post treatment, while Treg and ILC2 populations within this tissue remained suppressed throughout this time (Fig. 3L-P). Indeed, the mAT Th2 _RM_ population not only persisted, but continued to expand over time in the absence of infection (Fig. 3P).

### Resident Th2 cells and stromal cells activate each other during infection

When we examined mAT stromal cell activation in infected *Il4ra*^fl/fl^ *Cd4*-Cre mice, which lack Th2 cells, we found that cytokine and pCol1 production was diminished (Fig. 4A, B). There was also a failure of stromal cell activation when CD4^+^ T cells were depleted, an intervention that also resulted in the loss of resistance to infection (Fig. 4C-E, Fig. S7A-C). Th2 cells therefore play a critical role in mAT stromal cell activation during infection.

**Fig 4.**
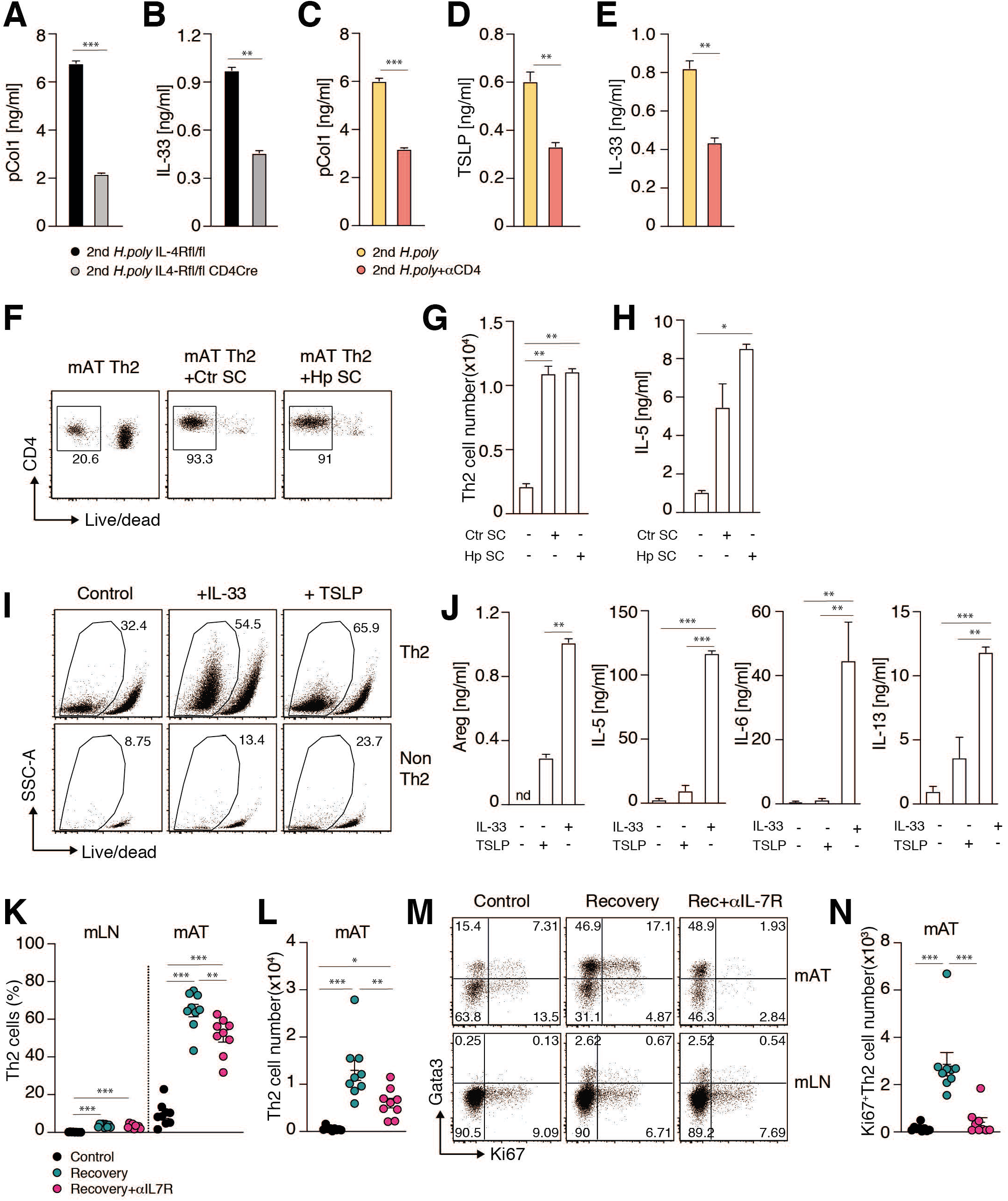
Interdependence of mAT Th2_RM_ and SC. (**A, B**) Secretion of pCol1 (A) and IL-33 (B) by isolated mAT SC of *Il4ra*^fl/fl^*Cd4*-*Cre* and *Il4ra*^fl/fl^ mice after secondary infection. (**C-E**) pCol1 (C), TSLP (D) and IL-33 (E) production from isolated mAT SC during secondary infection with anti-CD4 depletion where indicated. (**F, G)** Representative FACS plot (F) and cell number (G) of sorted Th2 cells after 4 days of co-culture with mAT SC isolated from control and infected mice as indicated. (**H)** IL-5 production by Th2 cells after 4 days of co-culture with mAT SC as in (F). (**I, J**). Th2 cells (CD4^+^ TCR*β*^+^ IL4-eGFP^+^ Foxp3RFP^-^) and non-Th2 CD4^+^ T cells (CD4^+^ TCR*β*^+^ IL4-eGFP^-^ Foxp3RFP^-^) were sorted from mAT and mLN of infected mice and cultured for 3 - 6 days with addition of IL-33 [50ng/ml] or TSLP [50ng/ml]. FACS plot of mAT Th2 and non-Th2 cells with frequencies of live cells after 6 days of culture (I). Levels of indicated cytokines in the supernatants of mAT Th2 cells after 3 days of culture (J). (**K-N**) TSLP signalling was blocked with anti-IL-7R*α* antibody treatment during recovery after primary infection. Frequencies (K) and numbers (L) of Th2 cells (gated on IL-33R^+^ GATA3^+^ among CD4^+^ TCR*β*^+^ T cells) in mAT or mLN as indicated. Representative FACS plot (M) and numbers (N) of Ki67^+^ Th2 cells in mAT. Dots represent BR (K, L, N), error bars represent SEM from BR (K, L, N) or TR (A-E, G, H, J). Data combined from 2 independent experiments (K, L, N) or representative of 2-3 experiments (A-E, G, H, J).

We asked whether SC reciprocally stimulate Th2_RM_ cells, by purifying and co-culturing these populations and measuring Th2 cell survival and cytokine production. We found both to be significantly enhanced in the presence of mAT SC from infected and naive mice (Fig. 4F-H). These effects were seen when mAT Th2_RM_ cells and mAT SC were cultured in a trans-well system, or when Th2_RM_ cells were cultured with stromal cell conditioned medium, indicating that soluble factor(s) from SC drive Th2_RM_ cell activation (Fig. S7D).

The expression of TSLPR and IL-33R on mAT Th2 cells, and the recognized relationship of TSLP and IL-33 with type 2 immunity, suggested that it could be these cytokines that are responsible for the observed effects. To examine this possibility, we sorted both Th2 cells (CD4^+^ Foxp3^-^IL-4^+^) and non-Th2 T cells (CD4^+^Foxp3^-^IL-4^-^) from mAT and mLN of infected animals and cultured them in the presence or absence of IL-33 and TSLP. We found that mAT Th2_RM_ cells were intrinsically more capable of surviving *in vitro* than non-Th2 cells or mLN Th2 cells, and that they proliferated extensively in the absence of added growth factor (Fig. 4I, Fig. S7E). This distinction was further enhanced by the addition of IL-33 or TSLP (Fig. 4I, Fig. S7E). Furthermore, these cytokines activated mAT Th2 cells to produce Areg, IL-5, IL-6, IL-13 and TGFβ_1_ in an antigen independent manner (Fig. 4J, Fig. S7F).

While IL-33-induced cytokine production more strongly than TSLP, TSLP had a comparable effect on mAT Th2_RM_ survival, suggesting that TSLP might be more important in the long-term maintenance of these cells. The TSLP receptor is a heterodimer of the Crlf2 and IL-7R*α*, the latter of which, when paired with *γ*_c_ (encode by *Il2rg*) is also a component of the *bona fide* IL-7R. Both TSLP and IL-7 receptors signal through STAT5 to promote T cell and ILC homeostasis (*21*). Th2_RM_ cells express *Crlf2, Il7ra* and *Il2rg* suggesting that they may use TSLP or IL-7 to establish their dominance in the mAT lymphoid niche (Fig. 2J, Fig. S8A-C). Antibody-mediated blockade of IL7R*α* for one week during post-infection recovery resulted in significant decreases in Th2 cells, and in Ki67^+^ Th2 cells in mAT, but not in mLN, confirming a requirement for TSLP/IL-7 signaling for Th2_RM_ cell proliferation and persistence (Fig. 4K-N). The decline in mAT Th2_RM_ cells caused by IL-7R*α* blockade also resulted in a decrease in mAT eosinophilia (Fig. S8D-F). Since we could not detect IL-7 in SC culture supernatants (*not shown*) and *Il7* was not strongly expressed in scRNAseq data (Fig. S8G), these results likely reflect the inhibition of TSLP-mediated effects. Together, our data point to the existence of a positive feedback loop in which Th2_RM_ cells in the mAT activate SC to secrete IL-33 and TSLP, which in turn promote mAT Th2_RM_ cell expansion, survival and cytokine production.

### Activated stromal multipotent progenitor cells accumulate and secrete collagen and immunostimulatory cytokines

To more fully explore the mAT stromal response to infection we re-clustered the scRNAseq stromal cell transcriptomes and identified 6 distinct cell groups (C0-C5, Fig. 5A, Fig. S9A, B). Using recently published transcription profiles to delineate adipocyte differentiation stages, we identified *Dpp4^+^ Pi16^+^* MPC (C2), intermediate uncommitted cells (C0), as well as *Fabp4^+^ Pparg^+^* committed preadipocytes (C1) (*14, 22, 23*). Other clusters were enriched in *CD9^+^* matrix fibroblasts (C3) (*24*) and *Ccl19+* immunofibroblasts (C4) (*25*) (Fig. 5A, Fig. S9A, B). By pseudotemporal ordering using Monocle (*26*) with MPC (C2) set as the origin, our data conformed with the preadipocyte to adipocyte differentiation model proposed by others (*14*) (Fig. 5B). A quantitative assessment of cluster sizes revealed an ∼25% increase in MPC (C2) and a decrease in intermediate uncommitted cells (C0), in infected versus control mice (Fig. 5C), suggesting a block in the differentiation of the MPC towards the adipocyte lineage.

**Fig 5.**
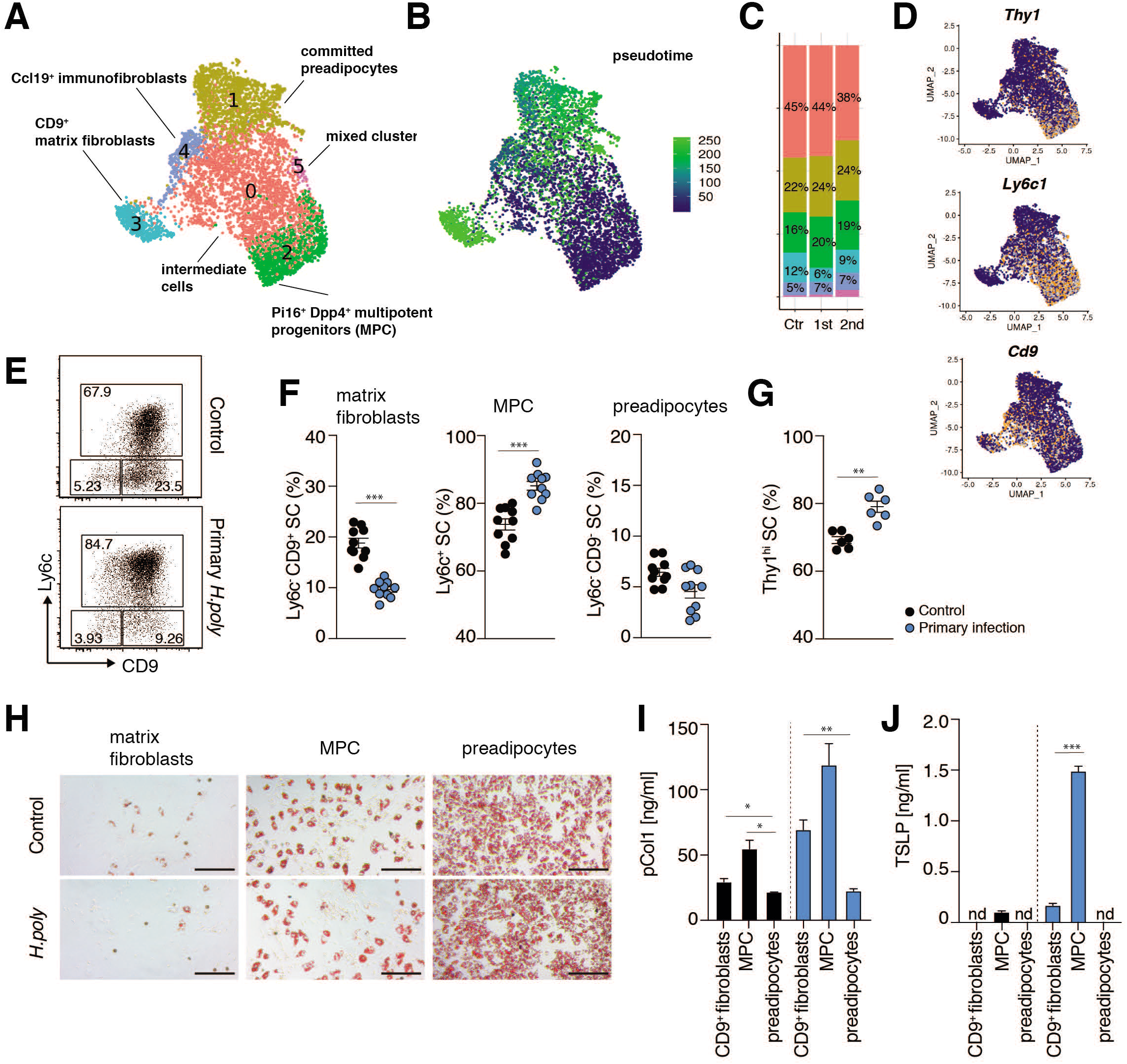
Expansion of the mAT *Dpp4^+^* MPC in *H. polygyrus* infection. (**A**) UMAP plot of 6059 mAT SC from control (2457 cells), primary infected (2640 cells) and secondary infected mice (962 cells). Unsupervised clustering distinguished 6 cell clusters (A); plots are colour-coded according to cell cluster. Identified cell population based on expression of following markers: committed preadipocytes (C1): *Icam1, Apoe, Lpl, Fabp4, Pparg*; pluripotent progenitors (C2): *Dpp4, Anxa3, Cd55, Pi16, Dpt*; CD9^+^ profibrotic cells (3): *Cd9, Wnt6, Eln, Mgp, Col1a1, Col15a1*; immunofibroblasts (C4): *Cd9, Ccl19*. (**B)** UMAP plot as in A, showing pseudotemporal ordering of cells, setting origin at the centre of C2 (multipotent progenitor cells, MPC). (**C**) Bar graphs represent contribution of each cluster as in (A) to a total cell pool, split by experimental condition. (**D**) UMAP plots indicating expression of selected genes. (**E, F**) Representative FACS plots (E) and quantification (F) of mAT SC populations from control and *H. polygyrus* infected mice: Ly6c^+^ progenitors, Ly6c^-^ CD9^+^ matrix fibroblasts and Ly6c^-^ CD9^-^ preadipocytes (gated on live CD45^-^ CD31^-^ PDGFR*α*^+^ cells). (**G**) Frequencies of Thy1^hi^ mAT SC in control and *H. polygyrus* infected mice (gated on live CD45^-^ CD31^-^ PDGFR*α*^+^ cells). (**H**). mAT SC were sorted into Ly6c^+^ progenitors, Ly6c^-^ CD9^+^ matrix fibroblasts and Ly6c^-^ CD9^-^ preadipocytes and subjected to adipogenic differentiation (see Methods). Representative images on day 6 showing accumulation of lipid droplets. Scale bar 200μm. **(I, J)** pCol1 (I) and TSLP (J) levels measured in supernatants from overnight culture of cells sorted as in (H). Dots represent BR (F, G), error bars represent SEM from BR (F, G) or TR (I, J). Data combined from 2 independent experiments (F), representative of two independent experiments (G, H-J) or from one experiment (A-D).

We used flow cytometry to address this further, gating on CD45^-^ CD31^-^ Sca1^+^ PDGFR*α*^+^ SC and using antibodies against surface molecules identified previously (*14, 23*) and in our data as marks of particular clusters, to quantitate MPC, which more strongly express *Ly6c* and *Thy1* (C2), versus fibroblast clusters, which more strongly express *Cd9*, or CD9^lo^Ly6c ^lo^ committed preadipocytes (C1) (Fig. 5D). Infection led to increased frequencies of MPC and reduced frequencies of committed preadipocytes (Fig. 5E-G). To verify functional differences between identified stromal subpopulations we sorted them based on Ly6c and CD9 expression (Fig. 5E) and asked which had the potential to become adipocytes under adipogenic culture conditions (*14*). We found that the committed preadipocytes had the highest adipogenic potential, evident by extensive lipid droplet development, while MPC showed intermediate adipogenic potential. The matrix fibroblast subpopulation contained few cells that were able to differentiate into mature adipocytes (Fig. 5H, Fig. S9C). Infection did not affect the inherent ability of cells within the different stromal subpopulations to differentiate into mature adipocytes in culture (Fig. 5H, Fig. S9C). These data confirmed the functional relatedness of the clusters identified in our study to previous descriptions of adipocyte differentiation (*14, 23, 24*).

We asked whether infection induced stroma cell remodeling could be attributed to a particular subpopulation of mAT SC. We found that pCol1 was produced by sorted matrix fibroblasts and MPC, and that both of these populations produced more pCol1 when sorted from infected mice (Fig. 5I). However, the MPC made more pCol1 than did the matrix fibroblasts, despite indications from the scRNAseq data that the opposite would be the case (discussed below). By comparison, committed preadipocytes from infected mice made little pCol1 (Fig. 5I). Additionally, MPC were the major source of TSLP, production of which was greatly increased as a result of infection (Fig. 5J).

Of relevance, given the expression of the IL-33R by mAT Th2_RM_ cells, and the effects of this cytokine on these cells, MPC expressed *Il33* more strongly than any of the other stromal populations, in line with previous reports (*7, 8*) (Fig. S9D). They also expressed the chemokine encoding genes *Ccl2*, the ligand for CCR2, which is expressed on Th2_RM_ cells, and *Ccl11,* which encodes an eosinophil attractant (Fig. S9D), as well as genes encoding ECM components and modifying enzymes, including *Fn, Postn, Ugdh, Pcolce2*, and *pCol1a2* (Fig. S9E, F), although expression of the latter, as well as *pCol1a2*, *pCol1a1, pCol3a1* and *pCol6a1*and *Eln* was strongest in matrix fibroblasts in the stroma (Fig. S9F).

Together, our findings support the view that there is an infection-associated shift in the mAT stromal cell population structure such that MPC accumulate in the tissue and that this is coincident with a reduction in organ size and fat storage. The MPC express genes which indicate that they are able to respond to, recruit and stimulate Th2_RM_ cells, and attract eosinophils, while at the same time modulating the tissue ECM.

### Activated stromal cells are critical for host protective immunity to infection

Our data indicated that mAT Th2_RM_ cells are able to interact with SC, in a manner that allows reciprocal functional regulation in the context of an intestinal infection. We used Cell Phone (*27*) to specifically interrogate potential interactions between Th2_RM_ cells and MPC, the stromal cell population which showed the most dynamic changes during infection. The results of this analysis emphasized interactions involving the products of *Tslpr* and *Tslp*, as well as the chemokine receptor *Ccr2* with *Ccl2* and *Ccl11* and interactions involving integrin complexes expressed by mAT Th2_RM_. It also highlighted interactions between Th2_RM_-derived TGFβ_1_ and its receptors and the possibility of *Areg-EGFR* crosstalk (Fig. 6A).

**Fig 6.**
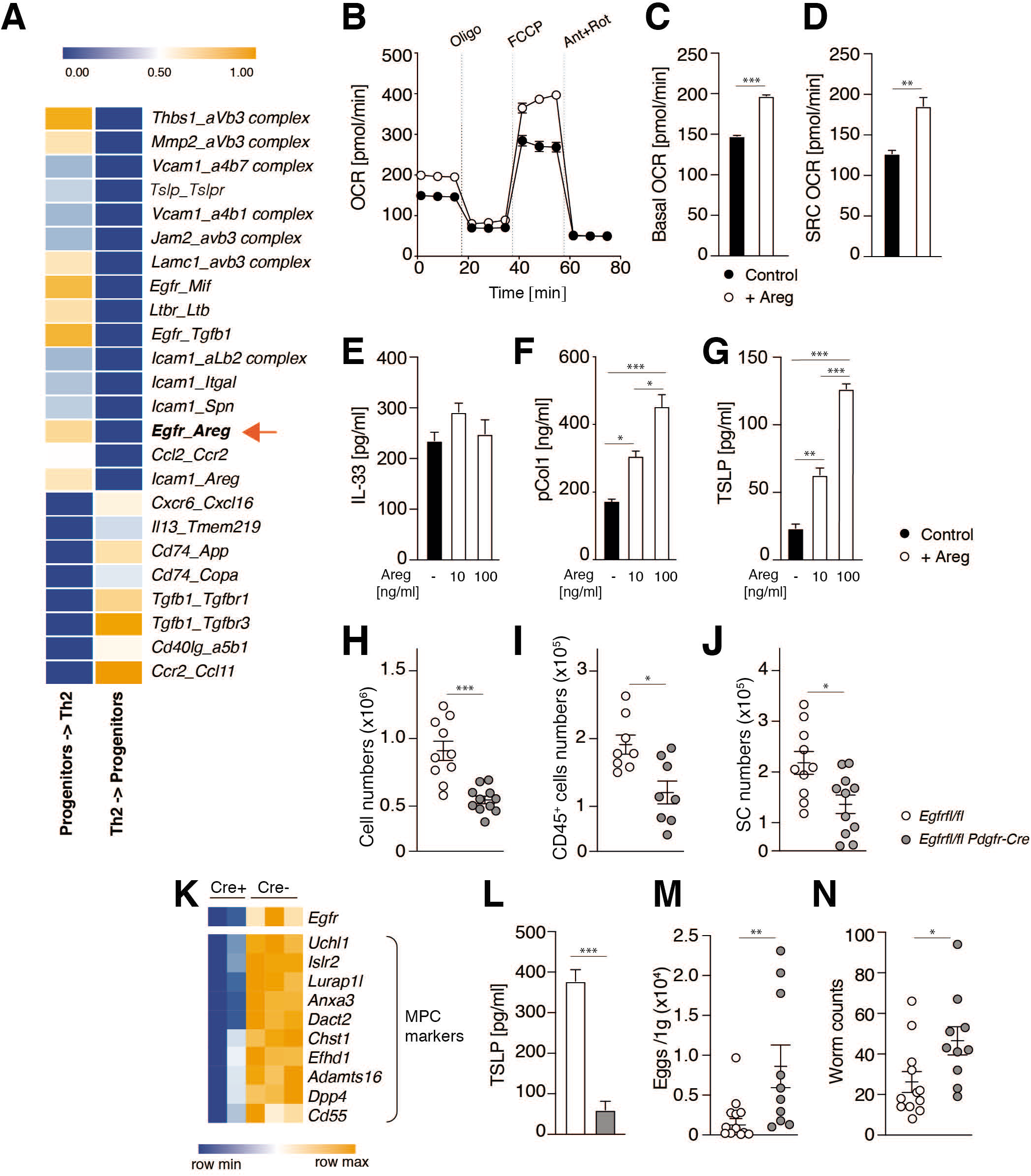
Areg supports mitochondrial respiration and TSLP production by mAT SC. (**A**) CellPhone analysis of ligand-receptor pairs between mAT Th2 cells and MPC. (**B-D**) mAT SC were subjected to 3 days of adipogenic differentiation in the presence of Areg [100ng/ml] where indicated. OCR was measured at baseline and after Oligomycin (Oligo), FCCP and Rotenone/Antimycin (Rot/Ant) injections (B). Baseline OCR (C) and SRC (difference between response to FCCP and basal OCR) (D). (**E-G**) mAT SC were subjected to 4-5 days of adipogenic differentiation in the presence of Areg as indicated. Levels of IL-33 (E), pCol1 (F) and TSLP (G) were measured in the supernatants. (**H-J**) Total SVF cell counts (H), immune cell counts (gated on live CD45^+^ CD31^-^ cells) (I) and SC counts (gated on live CD45^-^ CD31^-^ PDGFR*α*^+^) (J) in mAT of *Egfr*^fl/fl^ and *Egfr*^fl/fl^-*Pdgfra*-Cre mice. (**K**) Expression pattern of MPC markers in RNAseq dataset from mAT SC isolated from *H. polygyrus* infected *Egfr*^fl/fl^ and *Egfr*^fl/fl^-*Pdgfra*-Cre mice (sorted CD31^-^ CD45^-^ PDGFR*α*^+^). (**L**). Purified mAT SC from *H. polygyrus* infected *Egfr*^fl/fl^ and *Egfr*^fl/fl^-*Pdgfra*-Cre mice were cultured overnight and TSLP levels were measured in the supernatants. (**M, N**) Eggs in the caecum content (M) and worms in the small intestine (N) were enumerated in *Egfr*^fl/fl^ and *Egfr*^fl/fl^-*Pdgfra*-Cre mice with primary infection. Dots represents BR (H-J, L-N), error bars represent SEM from BR (B-D, F, H-J, L-N) or TR (E, G, L). Data combined from 2 independent experiments (H-J, M, N), representative of two to three independent experiments (B-G) or from one experiment (A, K, L).

We were intrigued by the role of Areg, since relatively little is known about its effects on AT biology. We began by broadly examining the effects of Areg on mAT SC in an *ex vivo* culture system. We found that mAT SC cultured in the presence of Areg had higher basal OCR and spare respiratory capacity (SRC) (*28*), indicating marked effects on metabolism (Fig. 6B-D). While the presence of Areg did not affect the secretion of IL-33 by mAT SC, it promoted the production of pCol1 and TSLP (Fig. 6E-G). Thus, Areg stimulated mAT SC from naïve mice shared metabolic and functional characteristics with activated mAT SC from infected mice. We then examined the effects of Areg on mAT by directly injecting naïve mice with this cytokine. We found increased numbers of immune and SC (Fig. S10A-C), which is consistent with the observed effects of infection. We then generated mice which lack EGFR on SC using *Pdgfra-*Cre (*Egfr*^fl/fl^-*Pdgfra*-Cre mice); *Pdgfra* is broadly expressed in SC and anti-PDGFR*α* was used here to sort SC from mAT. Consistent with a role for Areg in modulating mAT SVF cellularity, we found fewer SC and immune cells in the mAT of *Egfr*^fl/fl^-*Pdgfra*-Cre mice that in control mice (Fig. 6H-J). To gain insight into the role of EGFR in mAT SC biology during infection, we performed RNAseq of mAT SC isolated from infected *Egfrfl/fl-Pdgfra-*Cre and control mice. Lack of EGFR resulted in reduced expression of several genes characteristic of AT MPC, including *Dpp4, Anxa3* and *Cd55* (*14, 29*) (Fig. 6K), suggesting that Areg signaling through EGFR plays a role in maintaining the identity of the progenitor population during infection. We also found that mAT SC from infected *Egfr*^fl/fl^-*Pdgfra*-Cre mice produced less TSLP *ex vivo* than did SC from infected controls (Fig. 6L). Lastly, *Egfr*^fl/fl^-*Pdgfra*-Cre mice were more susceptible to *H. polygyrus* infection compared to control mice (Fig. 6M, N). Together, these data indicate that EGFR expression on SC regulates adipocyte lineage differentiation and immunity to *H. polygyrus*.

## Discussion

Here we show that the mAT response to an enteric parasitic infection is marked by coordinated, interactional changes in the immune and stromal compartments. Most strikingly, a population of Th2_RM_ cells expands to dominate the mAT lymphocyte niche, and in response to signals from SC, provides activating cytokines that drive the functional reprograming of the stroma. These effects encompass major changes in anabolic metabolism, gene expression and developmental trajectories that result in MPC becoming ECM-, cytokine- and chemokine-producing cells. The fact that these changes are exaggerated after exposure to secondary infection indicates that the tissue retains memory of the primary event. Understanding how Th2_RM_ cells regulate tissue biology through effects on tissue SC holds promise for the development of targeted tissue-regenerative therapeutics.

Pathways of adipocyte SC differentiation were defined in detail recently, but how these are modulated by physiological perturbations has so far been studied mostly in the context of obesity (*8, 14, 23*). We found that the Th2 response associated with *H. polygyrus* infection had a significant impact on the composition of mAT SC populations, characterized by a shift towards *Dpp4^+^ Pi16^+^* MPC, which became the primary producers of cytokines and collagen within the stroma. Our results are reminiscent of the situation in chronic rhinosinusitis, where type 2 inflammation causes a shift into a non-committed state in epithelial cells at sites of polyp formation (*30*). *Dpp4^+^ Pi16^+^* stromal progenitors were recently identified across different tissues in mice and humans, as a universal reservoir population containing cells capable of giving rise to differentiated fibroblast subsets (*29*). Further work is required to fully understand the population structure of progenitor cells in the context of infection, but it is of interest that apparent plasticity between adipocyte and fibroblast lineages during tissue damage has been noted (*31, 32*). Taken together the findings suggest the possibility that expansion and activation of the universal reservoir SC population might be a hallmark of the physiological response to type 2 inflammation that is shared across tissues. We speculate that the benefit of such a response is linked to the plasticity of SC in the universal reservoir population to assume new supportive functions in response to signals received from the immune system. In the case of infection with *H. polygyrus*, we believe that the accumulation of the MPC population effectively stalls differentiation into committed preadipocytes, a process that would be expected to result in the reduction in AT mass associated with infection, and which may explain previous reports that *H. polygyrus* infection can prevent obesity (*11*).

mAT activation in infected mice shares some features, including the activation of SC to make Col1A1, with the creeping fibrotic mAT of Crohns disease, that serves to prevent the systemic spread of intestinal bacteria which translocate across the gut wall due to loss of epithelial integrity associated with the disease (*33*). *H. polygyrus* are not thought to penetrate the serosal surface, but based on the creeping mAT model, we speculate that increased collagen deposition within the mAT may reflect a defensive process aimed at increasing the strength and resilience of the intestine and its associated vasculature to minimize the possibility and consequences of perforation. Despite the presence within the mAT of *CD9^+^* matrix fibroblasts, which strongly expressed collagen genes, the MPC were the main source of pCol1 protein during infection. The mechanical stiffness of the ECM is capable of influencing stem cell fate determination (*34*), so it is feasible that changes in ECM observed here could contribute to the accumulation of MPC during infection. Perhaps related to this, MPC differentiation can also be restrained by TGFβ_1_ (*14*), and in this context it is notable that mAT Th2_RM_ cells produced this cytokine. Both TGFβ_1_ and the Th2 cell cytokine IL-13 are strongly implicated in fibrotic disease (*35*), although infection-associated changes in ECM in mAT did not develop into persistent fibrotic remodeling during infection (data not shown), indicating that inflammatory and healing responses in mAT are well controlled in this setting.

Persistence of a large population of mAT Th2_RM_ cells almost a year post-clearance of infection was striking and consistent with reports of the longevity of lung Th2_RM_ cells (*36*). Th2_RM_ cells within mAT expressed *Arg1, Nmur1* and *Calca*, which have previously been considered to be primarily expressed by ILC2 (*37–40*). This pattern of gene expression, together with their ability to become activated in an antigen-independent manner, supports the view that innate reprograming of Th2 cells is an integral part of terminal differentiation driven by exposure to tissue-derived cytokine such as TSLP and IL-33 (*17, 41, 42*). We speculate that immunologic remodeling of mAT with adaptive immune cells that have acquired the ability to become activated in response to innate signals may influence the susceptibility of this tissue to other insults, such as the establishment of cancer metastases, or infection by pathogens that require Th1 or Th17 responses for resolution.

In addition to classic type 2 cytokines, mAT Th2_RM_ cells also produced the growth factor Areg. The fact that Areg can release TGFβ_1_ from latent TGFβ_1_ through integrin-*α*_v_ activation (*43*) indicates that polyfunctional Th2_RM_ cells capable of making both cytokines may be particularly potent sources of active TGFβ_1_. Relatively little is known of roles for Areg and EGFR signaling in AT physiology. We observed that Areg can drive TSLP production by mAT SC. This crosstalk between Areg and TSLP production could stabilize the lymphoid niche within the tissue and therefore have implications for the persistence of Th2_RM_ cells in mAT. Deletion of *Egfr* also emphasized the importance of Areg signaling for modulating SVF cellularity in mAT, and for protective immunity against an enteric infection. While our experiments did not allow identification of mAT SC as those critical for immunity, they nevertheless support an emerging view of immune cell driven EGFR signaling in SC providing a critical component of tissue immunity against infection.

Our findings on mAT fit with the growing realization that AT can provide help to tissues fighting infection or recovering from wounding (*13, 44–46*). These findings warrant broader consideration of the function of AT during disease. The extent to which changes in populations of resident immune cells affect the helper activity of AT has been unclear, but our findings indicate that this may be of major significance since immune cells and AT SC have evolved powerful dynamic mechanisms for reciprocal activation and regulation.

## Acknowledgments

The authors thank Drs. Mark Wilson, Asifa Akhtar, Maximilian Seidl, Rick Maizels, Peter Murray, Sabine Eming, Axel Roers and David Vöhringer for reagents and helpful discussions, the core facilities at the MPI-IE and at the Institute of Clinical Pathology, University of Freiburg, for their support, and Andrea Quintana, John Sutherland and Fabian Haessler for help with animal colonies.

## Funding

National Institutes of Health grant AI 110481 (EJP)

German Research Foundation grant DFG FOR 2599 (EJP, AMK)

German Research Foundation grant under Germany’s Excellence Strategy CIBSS EXC-2189 Project ID 390939984 (DG)

German Research Foundation grant SPP1937 GR4980/1-1 (DG) Alexander von Humboldt Fellowship Foundation (AMK, MV, FB, JC)

CAPES/Alexander von Humboldt Fellowship Foundation grant 88881.136065/2017-01 (AC) Marie Skłodowska-Curie action Individual fellowship MSCA-IF (FB, JC) European Research Council Advanced grant ERC-2015-AdG TNT-Tumors 694883 (MS)

European Union’s Horizon 2020 research and innovation program under the Marie Skłodowska-Curie grant agreement No. 766214 Meta-Can (MS)

Max Planck Society

## Author contributions

Conceptualization: AMK, DES, JAP, ELP, EJP

Methodology: DES, PZ, NR, JEH, DG

Investigation: AMK, AH, KMG, PZ, LD, RK, GC, JDC, AC, MV, FB, JC

Visualization: AMK, FB, DES, PZ

Funding acquisition: ELP, EJP

Supervision: EJP

Resources: JFU, MS Writing: AMK, EJP

## Methods

### Mice and *H. polygyrus* infection model

We used C57BL/6J, (JAX:000664), *Cd4-*Cre (JAX: 022071), *Pdgfra-*Cre (JAX:013148), *Il4*eGFP *Foxp3*RFP *Il10*Bit (generated by crossing *Il4*^tm1Lky(47)^ (*47*), *Foxp3*^tm1Flv^ (*48*) and Tg(Il10-Thy1a) (*49*) mice, kindly provided by M. Wilson), *Il4ra^tm2Fbb^* (IL-4Rfl/fl) (*50*), kindly provided by F. Brombacher, and *Egfr^tm1Dwt^* (EGFRfl/fl) (*51*) mice. All mice were maintained in specific-pathogen-free conditions in the animal facility of the Max Planck Institute for Immunobiology and Epigenetics (Freiburg, Germany), and all corresponding animal protocols were approved by the animal care committee of the Regierungspraesidium Freiburg. For *H. polygyrus* infection, mice were gavaged with 200 infectious L3 stage larvae in PBS. For primary infection, mice were left for 11-14 days before being sacrificed or treated with the anthelminthic pyrantel pamoate (1mg/mouse). For secondary infection, mice were infected at 5 weeks post treatment, and sacrificed 11-14 days later. For CD4^+^ T cell depletion, mice were infected and treated, and 5 weeks later injected with anti-CD4 monoclonal antibody (mAb, clone GK1.5, BioXCell, 500 μg /mouse, i.p. per injection) one day before secondary infection, and then again at day 6 after infection. Mice were sacrificed on day 11 of secondary infection. For IL-7R*α* blockade, mice were infected and treated, and injected with anti-IL-7R*α* mAb (clone A7R34, BioXCell, 500 μg/mouse, i.p. per injection) on days 3 and 9 post treatment, and sacrificed on day 12 post treatment. For treatment with FTY720 (Enzo, BML-SL233-0005), mice were infected and treated, and 5 weeks later injected with FTY720 every second day, starting from one day before secondary infection (6 injections in total, 10 μg/mouse, i.p. per injection). Mice were sacrificed on day 11 of secondary infection. For Areg (R&D System, 989-AR) treatment, naïve mice were injected 3 times with 10 μg/mouse Areg i.p. on day 0, 3, and 6 and were sacrificed on day 9. In all experiments mice were age and sex matched. Both females and males were used. For the magnetic resonance imaging (MRI), an EchoMRI 3-in-1 Body Composition Analyzer (EchoMRI) was used according to the manufacturer’s instructions.

### *H. polygyrus* egg and adult worm counts

Small intestines were removed, opened longitudinally, and placed into a mesh cloth on top of a 50 ml tube filled with PBS for 3-4 h in a 37°C water batch. Parasites dropped through the mesh into the tube and were recovered for counting on a dissecting microscope. Parasite eggs in caecal contents collected from individual mice were collected and enumerated by floatation on sodium chloride and counted under a microscope.

### Isolation of cells from mAT and SI LP

For isolation of cells from mAT, mice were euthanised and transcardially perfused with 10 ml of ice-cold PBS. mAT was surgically separated from mLN, intestine and omentum, minced and digested in 3 ml of low glucose DMEM containing 25 mM HEPES, 1% low fatty acid BSA, 2 mM L-glutamine (L-glut), 100 U/ml Penicillin/Streptomycin (P/S), with the addition of Liberase TL (0.2 mg/ml, Roche) and DNase I (0.25 mg/ml, Roche) for 30-40 min at 37°C on a rotator. After digestion, 2 ml DMEM containing 2 mM EDTA was added and the suspension was filtered through 70 µm strainers. Cells in SVF were separated from the adipocyte layer by centrifugation. For isolation of cells from SI LP, SI were opened longitudinally, cleaned with PBS, cut into 2 cm pieces and placed in RPMI containing 3% FBS, 25 mM HEPES, 2 mM L-glut, 100 U/ml P/S, 1 mM DTT and agitated for 25 min, after which samples were washed 3-4 times with RPMI containing 2 mM EDTA. Intestines were digested in RPMI containing 25 mM HEPES, 2 mM L-glut, 100 U/ml P/S, Liberase TL (0.1 mg/ml, Roche), DNase I (50 µg/ml, Roche) for 30 min at 37°C with agitation. Cell suspensions were filtered through 70 µm strainers, centrifuged and resuspended in 3ml P30 solution (30% Percoll, GE, Amersham, UK). Overlaying 3 ml P75 with 4 ml P40 and then 3 ml P30 containing SI LP leukocytes created a three-layered discontinuous gradient. Gradients were centrifuged (1800 rpm/ 680g, 20°C, 20 min) and enriched leukocytes isolated from the P40/P75 interface.

### SC cultures

SC were isolated from SVF using the Adipose Tissue Progenitor Isolation Kit (Miltenyi Biotec, #130-106-639). Isolated SC were plated at 2-4×10^5^ cells/well in 48 well plates and cultured overnight at 37°C, 5% CO_2_ in cRPMI medium (RPMI containing 10% FBS, 25 mM HEPES, 2 mM L-glut, 100 U/ml P/S, cRPMI). Supernatants were then collected and cytokines or pCol1 measured by ELISA. In experiments where effects of Areg on SC biology were assessed over longer culture periods, adipogenic factors were added to mimic adipose tissue conditions. SC from mAT of naïve mice were seeded at near confluency and cultured in complete DMEM/F12 medium (DMEM/F12, 10% FBS, 1 mM sodium pyruvate, 2 mM L-glut, 100 U/ml P/S) supplemented with dexamethasone (Sigma, #D-1756, 0.5 μM), 3-isobutyl-1-methylxanine (IBMX, Sigma, #I-5879, 0.5 mM), insulin (Sigma, #I-5523, 1.7 μM) and Rosiglitazone (Sigma, #R2408, 1μM). Medium was changed every second day and on day 4 was replaced with adipogenic medium containing only insulin. Where indicated, cells were cultured with Areg (R&D Systems, #989-AR-100, 10 or 100 ng/ml) or heptelidic acid (Adipogen, #AG-CV2-0118-M00, 10 μM).

In experiments where the adipogenic potential of mAT SC subpopulations was assessed, cells were sorted according to CD9 and Ly6c expression and plated at equivalent cell numbers in complete DMEM/F12 medium for 24 h, after which the medium was supplemented with minimal adipogenic medium: 1 μM dexamethasone (Sigma, #D-1756), 3-isobutyl-1-methylxanine (IBMX, Sigma, #I-5879, 0.5 mM), insulin (Sigma, #I-5523, 20 nM) and Rosiglitazone (Sigma, #R2408, 1μM). Medium was changed every second day of the culture and from day 4 adipogenic medium was supplemented only with 20 nM insulin. Lipid droplets were visualized on day 7 of the culture using Oil Red solution (Sigma, 01391): briefly, cells were fixed with 10% NBF (Sigma) for 30 min at RT, washed with H_2_O and incubated with 60% Isopropanol/ H_2_O for 5 min, followed by staining with 60% Oil Red/ H_2_O solution for 20 min. Cell were washed with H_2_O before microscopy. For semi-quantification, cells were additionally washed with 60% Isopropanol/ H_2_O, after which Oil Red was released from the cells by incubation with 100% Isopropanol for 5 min. Absorbance was read at 492 nm.

### Isolation and culture of mAT and mLN T cells

Triple reporter (*Il4*^tm1Lky^ *Foxp3*^tm1Flv^ Tg(Il10-Thy1a) mice were infected with *H. polygyrus* as described above and Th2 cells (TCR*β*^+^ CD4^+^ IL4-eGFP^+^ Foxp3-RFP^-^) and non-Th2 effector T cells (CD45^+^ TCR*β*^+^ CD4^+^ IL4-eGFP^-^ Foxp3-RFP^-^) were FACS-sorted, plated at 1-2×10^4^ cells/well in 96 U-bottom plate and cultured for 3 – 6 days in cRPMI medium in the presence of IL-33 (R&D Systems, #3626, 50 ng/ml) or TSLP (R&D Systems, #555-TS, 50ng/ml) where indicated. Cells were then enumerated and supernatants were collected for cytokine measurements. In co-culture experiments, SC from control or *H. polygyrus* infected mice were purified as described above and plated at 5×10^4^/well in flat bottom 96-well plates, or at 1.7×10^5^/well in the lower chamber of a 96 trans-well plate (Corning, 3381, 0.4μm polycarbonate membrane). One day later, sorted T cells were added to the upper chamber (1-2×10^4^ cells per well) and cultured for 3-4 days, after which cells were enumerated and supernatants collected for cytokine measurement.

### Histological staining

mAT and mLN tissue sections were fixed in buffered 10% formalin and paraffin-embedded. Sections were then cut and stained with hematoxylin and eosin. Images were acquired using a Zeiss Axio Imager Apotome microscope.

### Immunofluorescence staining and microscopy

mAT samples were fixed for 60 min on ice in 10% NBF (Sigma), permeabilized in 1% Triton X-100 in PBS (Sigma) for 30 min at room temperature (RT) and blocked for 30 min at RT with 2.5% BSA, 0.5% Triton X-100 in PBS. mAT samples were incubated with primary antibodies, anti-CD3 (Abcam, ab5690, rabbit polyclonal), anti-Perilipin-1 (Abcam, ab61682, goat polyclonal), anti-PDGFRa (R&D System AF1062, goat polyclonal) and GATA3-e660 (Invitrogen, mAb clone TWAJ) in 0.5% BSA, 0.5% Triton X-100 in PBS for 60 min at RT followed by three washes with 0.5% BSA, 0.5% Triton X-100 in PBS. After washing, samples were incubated with secondary antibodies conjugated with Alexa Fluor 488 and/or Alexa Fluor 568 (Thermofisher) in 0.5% BSA, 0.5% Triton X-100 in PBS for 60 min. All antibodies were used at 1:100 dilution. Nuclei were stained with Hoechst 33342 (Thermofisher, 2 μg/ml) in 0.5% BSA, 0.5% Triton X-100 in PBS for 30 min. Samples were mounted with ProLong® Diamond Antifade Mountant (Thermofisher). Confocal images were acquired using a Zeiss spinning disk confocal microscope equipped with a Photometrics Prime BSI camera and Apochromat objectives.

### Flow Cytometry

For analysis of intracellular cytokine production cells were re-stimulated for 4 hours at 37°C in RPMI media supplemented with 10% FCS, 2 mM L-glut, 100 U/ml P/S with 0.1 μg/ml Phorbol 12-myristate 13-acetate (PMA), 1 μg/ml Ionomycin and 10 μg/ml Brefeldin A. Cells were surface stained with mAbs diluted in PBS/0.1% BSA and Fc-block (Biolegend) for 30 min on ice. Fixable Viability Dye (eBioscience) was added to allow the exclusion of dead cells from the analysis. The following fluorochrome-conjugated antibodies from Biolegend or BD Bioscience were used: CD4 (clone RM4-5), CD9 (clone MZ3), CD11b (clone M1/70), CD11c (clone N418), CD24 (clone M1/69), CD29 (clone HMb1-1), CD31 (clone 390), CD45 (clone 30-F11), PDGFRa (clone APA5), Ly6A (E13-161.7), Ly6C (clone HK1.4), Ly6G (clone 1A8), CD44 (clone IM7), CD49d (clone R1-2), CD62L (clone MEL-14), CD64 (clone X54-5/7.1), CD69 (clone H1.2F3), F4/80 (clone BM8), IL-7R (clone A7R34), SiglecF (clone E50-2440), TCR*β* (clone H57-597), Thy1 (clone 53-2.2). Additionally, biotinylated antibodies were used for the staining ST2 (IL-33R) (Mdb, clone DJ8) and TSLPR (Biolegend, 22H9). For intracellular staining, cells were fixed and permeabilized using the Foxp3/transcription factor staining kit (eBioscience), and incubated with the following fluorochrome-conjugated mAb: GATA3 (clone TWAJ), Foxp3 (clone FJK-16s), IL-5 (clone TRFK5), IFN*γ* (XMG1.2), TGFB1-LAP (clone TW7-16B4), Ki67 (clone SolA15). Areg was detected with a biotinylated mAb (R&D Systems, BAF989). To identify ILC, a cocktail of biotinylated mAb against the following surface markers was used to allow gating on linage negative cells: NK1.1, CD19, CD5, F4/80, CD11c, Ter119, Cd3e, CD11b, LysG1, TCR*γδ*. Binding of biotinylated antibodies was detected using fluorochrome-conjugated streptavidin (Biolegend). Flow cytometry analyses were performed on a Fortessa flow cytometer (BD) and data were analyzed with FlowJo 9.9.4 software (BD).

### Cytokine and pCol1 measurements

Concentrations of Areg, IL-33, TSLP and pCol1 in cell culture supernatants were determined using ELISA kits (R&D System for Areg, IL-33 and TSLP, and Abcam for pCol1). Unless stated otherwise, data were normalized to 1×10^6^/ml cells for MSC supernatants and 1×10^5^/ml for T cell supernatants. For measuring IL-2, IL-4, IL-5, IL-6, IL-13 and TGFβ we used the LEGENDplex mouse Th Cytokine panel (Biolegend, #741044) and the Mouse/Rat Active/Total TGFβ assay (Biolegend, #740490).

### Metabolic Profiling

Oxygen consumption rates (OCR) and extracellular acidification rates (ECAR) were measured in XF media (non-buffered RPMI 1640 containing 25 mM glucose, 2mM L-glut, and 1 mM sodium pyruvate) under basal conditions and in response to 1 μM oligomycin, 1.5 μM fluoro-carbonyl cyanide phenylhydrazone (FCCP) and 100 nM rotenone + 1μM antimycin A using a 96 well XF or XFe Extracellular Flux Analyzer (EFA) (Seahorse Bioscience). 2×10^5^ freshly isolated mAT MSC were spun onto poly-D-lysine-coated seahorse 96 well plates and preincubated at 37°C for a minimum of 45 min in the absence of CO_2_. For the measurement of OCR and ECAR in MSC that were undergoing adipogenic differentiation, cells were culture in the seahorse 96 well plates in the presence of adipogenesis inducing factors as stated above for 3 days, after which medium was changed to XF media before the measurement. Glucose, glutamine and lactate concentrations MSC culture supernatants were measured using a Cedex Bio Analyzer (Roche).

### RNA sequencing

RNA isolations were done using Ambion’s RNAqueous Micro Kit (Cat 1931) according to manufacturer’s instructions and quantified using Qubit 2.0 (Thermo Fisher Scientific). Libraries were prepared using the TruSeq stranded mRNA kit (Illumina) and sequenced in a HISeq 3000 (Illumina) by the Deep-Sequencing Facility at the Max Planck Institute for Immunobiology and Epigenetics. Sequenced libraries were processed in an in-house developed RNA sequencing pipeline (*52*) that employs deepTools for quality control (*53*), cutadapt for trimming (DOI:10.14806/ej.17.1.200), STAR for mapping (*54*) and featureCounts to quantify mapped reads (*55*). Raw counts of mapped reads were processed in R (Lucent Techonologies) with DESeq2 (*56*) to determine differentially expressed genes and generate normalized read counts to visualize as heatmaps using Morpheus (Broad Institute). Pathway enrichment analysis was performed using in house application ‘Gene2Functions’ (https://github.com/maxplanck-ie/Genes2Functions).

### Single cell RNA sequencing: mAT SVF single cell sequencing

Single cell RNA sequencing of sorted mAT SVF cells was performed using a 10X Genomics Chromium Controller. Single cells were processed with GemCode Single Cell Platform using GemCode Gel Beads, Chip and Library Kits (v2) following the manufacturer’s protocol. Libraries were sequenced on HiSeq 3000 (Illumina). Samples were demultiplexed and aligned using Cell Ranger 2.2 (10X genomics) to genome build GRCm38 to obtain a raw read count matrix of barcodes corresponding to cells and features corresponding to detected genes. Read count matrices were processed, analyzed and visualized in R using Seurat v.3 (*57*) and Uniform Manifold Approximation and Projection (UMAP) (McInnes, L, Healy, J, UMAP: Uniform Manifold Approximation and Projection for Dimension Reduction, ArXiv e-prints 1802.03426, 2018) as a dimensionality reduction approach. Only cells with high quality transcriptomes (low % of mitochondrial RNA content and more than 200 detected genes) were used for data normalization and integration. Datasets were normalizes using a regularized negative binomial regression, and harmonized by identifying mutual nearest neighbors across datasets and applying canonical correlation analysis. Differentially expressed genes within each cluster and across conditions were determined with Seurat as those with a greater than 1.2-fold change and an adjusted p value of less than 0.05. Pseudotime was estimated using Monocle (*26*) setting pluripotent progenitor pre-adipocytes as the origin differentiation trajectories. Potential cell-to-cell interactions between selected populations were detected using CellPhone (*27*), where significant (p value < 0.05) means of the average expression levels of the interacting partner in the target cell types are presented as a heatmap. For the pathway enrichment analysis for the MSC clusters genes differentially expressed in each cluster compared to all other clusters where adjusted p value was lower than 0.01 and expression greater than 0.25 log FC were used to calculate pathway enrichment against GO terms in the biological process category. GO pathways where the sum of p values for each cluster was greater than 5.3 were then selected.

### Single cell RNA sequencing: CD4^+^ T cell single cell sequencing

Live CD45^+^ CD4^+^ TCR*β*^+^ T cells from mLN and mAT of control (2 mice per condition), primary infected (one mouse per condition) and secondary infected (one mouse per condition) and from SI LP of control and primary infected mice (2 mice per condition) were FACS sorted into 384-well plates containing lysis buffer (Herman et al., 2018). Plates were centrifuged for 5 min at 2200g at 4°C, snap-frozen in liquid nitrogen and stored at −80°C until processed. Single cell RNA sequencing was performed using the CEL-Seq2 method (*58*) with modifications as described (*59*). CD4^+^ T cells were sequenced on a HiSeq 2500 or HiSeq 3000 sequencing system (Illumina paired-end multiplexing run, high output mode) at a depth of ∼77,000 - ∼518,000 reads per cell.

Paired end reads were aligned to the transcriptome using bwa (version 0.6.2-r126) with default parameters (*60*). The transcriptome contained all gene models based on the mouse ENCODE VM9 release downloaded from the UCSC genome browser comprising 57,207 isoforms with 57,114 isoforms mapping to fully annotated chromosomes (1 to 19, X, Y, M). All isoforms of the same gene were merged to a single gene locus. Furthermore, gene loci overlapped by >75% were merged to larger gene groups. This procedure resulted in 34,111 gene groups. The loci for *Nmur1* was extended at the 3’ end to position 86,384,000 on the chromosome 1 (*39*). The right mate of each read pair was mapped to the ensemble of all gene loci in sense direction. Reads mapping to multiple loci were discarded. The left read contained the barcode information: The first six bases corresponded to the unique molecular identifier (UMI), followed by six bases representing the cell specific barcode. The remainder of the left read contained a polyT stretch. The left read was not used for quantification. For each cell barcode, the number of UMIs per transcript was counted and aggregated across all transcripts derived from the same gene locus. Based on binomial statistics, the number of observed UMIs was converted into transcript counts (*61*). The dataset was analyzed using RaceID3 (v0.1.3 and v0.1.6). Rescaling to 2,000 transcripts per cells was used for data normalization. Prior to filtering and normalization, mitochondrial genes were excluded and cells expressing >2% of *Kcnq1ot1* transcripts, identified as a marker of low-quality cells (*62*) were removed from analysis. This resulted in total 3431 CD4^+^ T cells with high quality transcriptomes and 26262 quantified genes across these cells. RaceID3 was run with the following parameters: mintotal= 2,000, minexpr=3, outminc=3, FSelect=TRUE, probthr=10^-3^, ccor = 0.65. In addition, we used imputing implemented in RaceID3 with knn=10. To remove batch effects and technical variability, we initialized the following sets of genes as CGenes: ensemble of all *Gm, RP, Hsp* and *A4300 genes, Malat1, Xist, Mid1 and Kcnq1ot1.* After initial clustering, cells within clusters with macrophage and plasma cell signature were removed and RaceID3 was rerun with the same parameters described above and *Lars2* was included in CGenes. In the final clustering step, VarID (*18*) was used with the distance matrix derived from RaceID3 clustering and the following pruning parameters: knn=10, alpha=1, FSelect=FALSE and the default density clustering parameters.

Due to recognized technical difficulties related to excessive mucus production, we were unable to isolate live SI cells from mice with secondary infection (*63*).

For the SVF scRNASeq analysis male mice were used, for the CD4^+^ T cell scRNASeq female mice were used.

### Fraction dot plot for clusters

Cluster-specific gene expression where color represents the Z score of the mean expression or log2 expression across clusters and dot size represents the fraction of cells in the cluster expressing the selected gene. For expression color code, log2 mean expression above for 4 or below −4 are replaced by 4 and −4, respectively. Z scores above 1 and −1 are replaced by 1 and −1, respectively.

### Quantification and Statistical Analysis

With the exception of scRNASeq and RNASeq dataset, statistical analysis for the remaining results were performed using Prism 7 software (GraphPad) and results are represented as Mean ±SEM. Comparisons for two groups were calculated using nonparametric Mann–Whitney test or unpaired two-tailed Student’s t tests. Comparisons of more than two groups were calculated using one-way ordinary unpaired ANOVA with Bonferroni’s multiple comparison tests. Designation of p-values were as follows: * *≤* 0.05; ** *≤* 0.01; *** *≤* 0.001.

For *in vivo* experiments, sample size was determined by power analysis and calculated required sample sizes were applied whenever possible. Selection of sample size for *in vitro* experiments was based on previous experience.

---

**Fig. S1.**
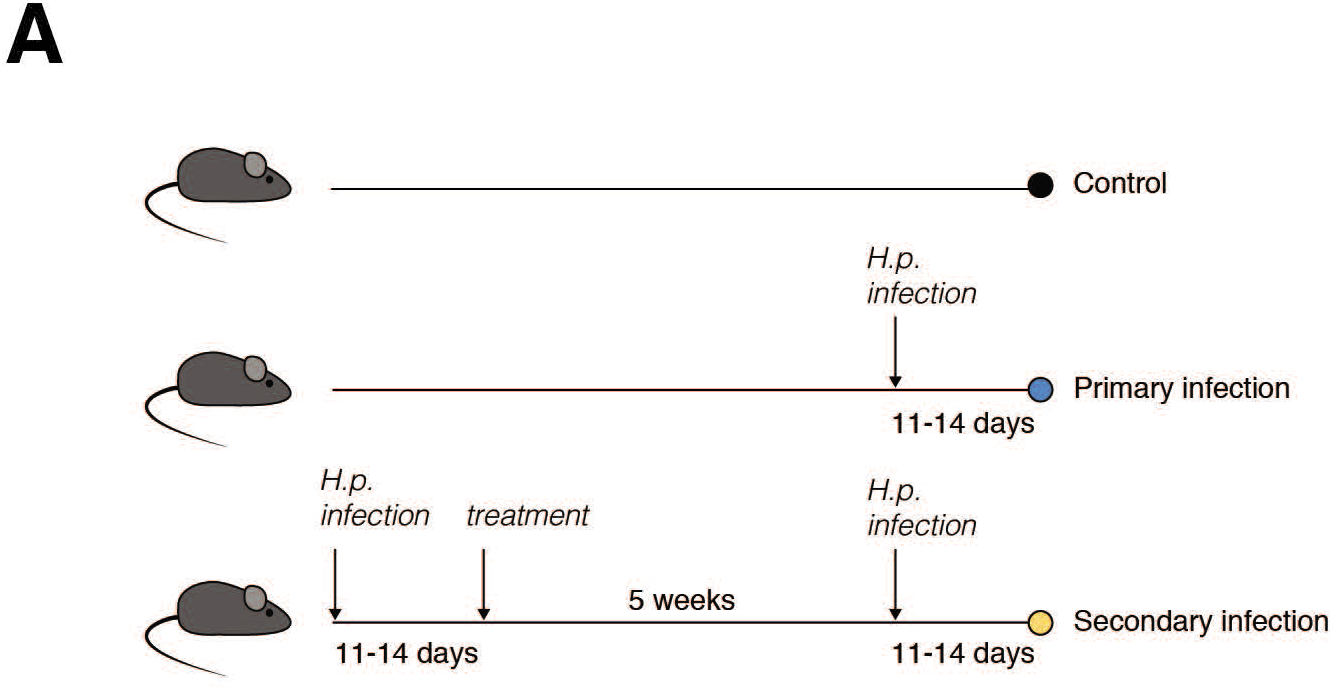
(**A**) Schematic representation of *H. polygyrus* infection model.

**Fig. S2.**
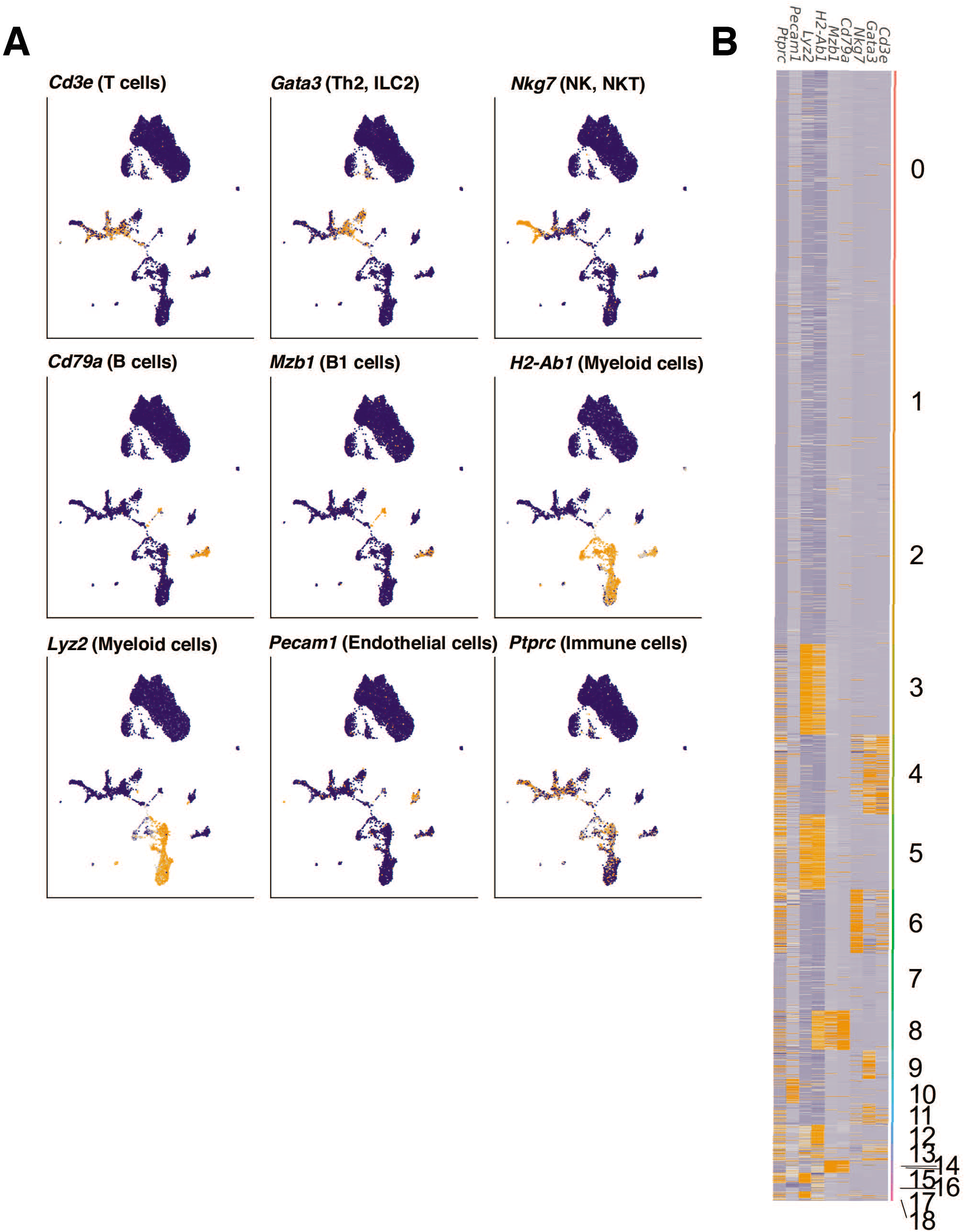
Marker gene expression pattern in SVF scRNASeq data from mAT. (**A**) UMAP plots of SVF cells isolated from mAT depots of control, primary infected and secondary infected mice as in Fig. 2A, expression of key genes used for identification of cell types is highlighted. (**B**) Heatmap showing selected key markers used for identification of cells in the clusters in the SVF scRNASeq dataset. Numbers indicate cluster identity. Data from one experiment.

**Fig. S3.**
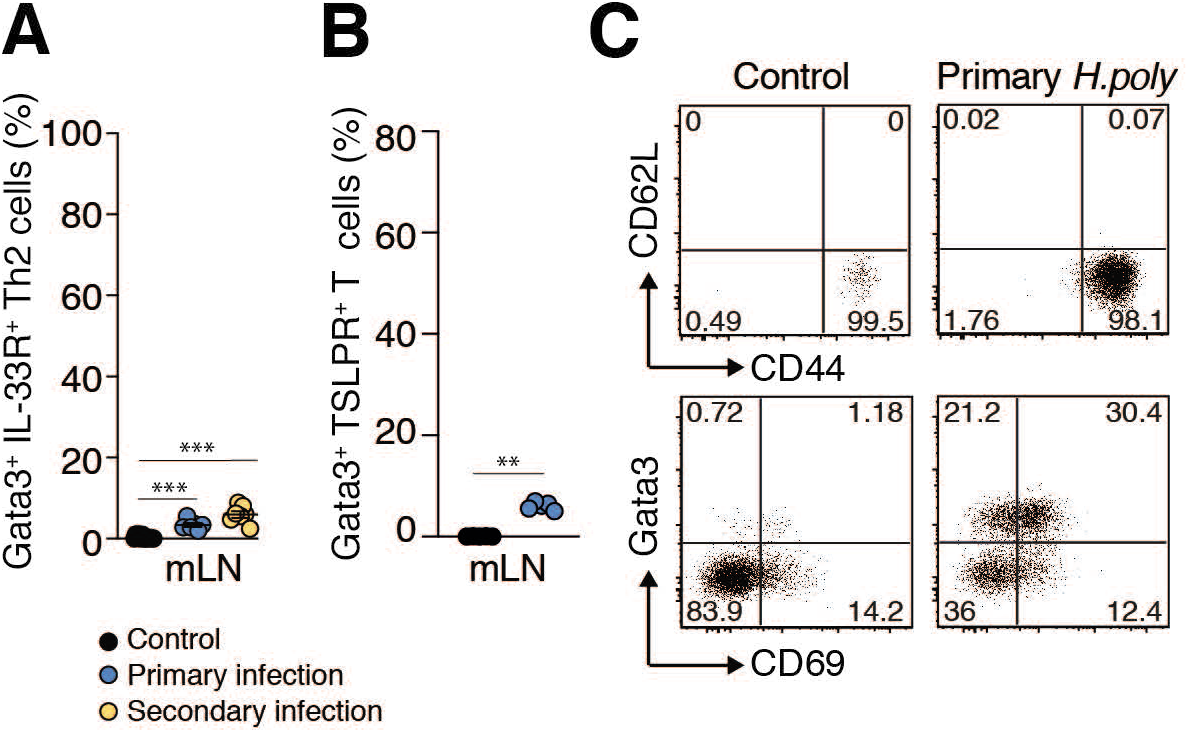
Characteristics of T cells in mLN and mAT of *H. polygyrus* infected mice. (**A, B**) GATA3^+^ IL-33R^+^ (A) and GATA3^+^ TSLPR^+^ (B) Th2 cells (gated on Foxp3^-^CD4^+^ TCRβ^+^ CD45^+^ live cells in lymphocyte gate) in mLN during *H. polygyrus* infection. (**C**) Representative FACS plot of CD44, CD62L and CD69 expression in mAT Th2 cells in control and *H. polygyrus* infected mice. For CD44, CD62L cells were gated on Foxp3^-^ GATA3^+^ TCRβ^+^ CD4^+^ CD45^+^ live cells in lymphocyte gate, for CD69 cells were gated on live Foxp3^-^ TCRβ^+^ CD4^+^ CD45^+^ cells in lymphocyte gate. Dot represents BR and error bars represent SEM from BR (A, B). Data combined from 2 independent experiments (A) or representative of 2 experiments (B, C).

**Fig. S4.**
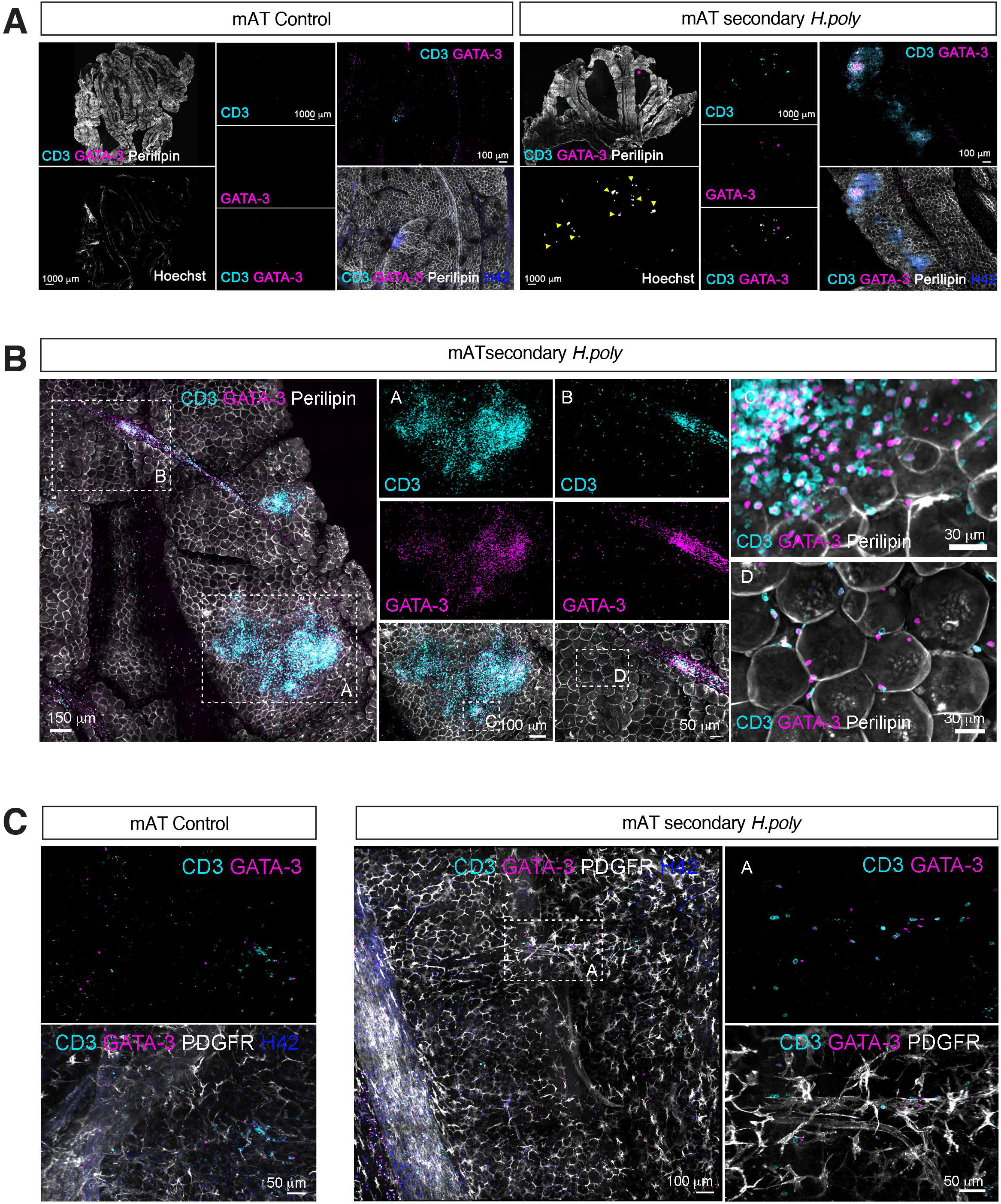
Localization of Th2_RM_ cells in mAT in *H. polygyrus* infection. (**A-C**) Whole mount images of mAT from control and secondary *H. polygyrus* infection stained for CD3, GATA3, Perilipin-1, PDGFR*α* and nuclear staining (Hoechst) as indicated. Yellow arrows indicate tertiary lymphoid structures with the mAT (A). Data from one experiment.

**Fig. S5.**
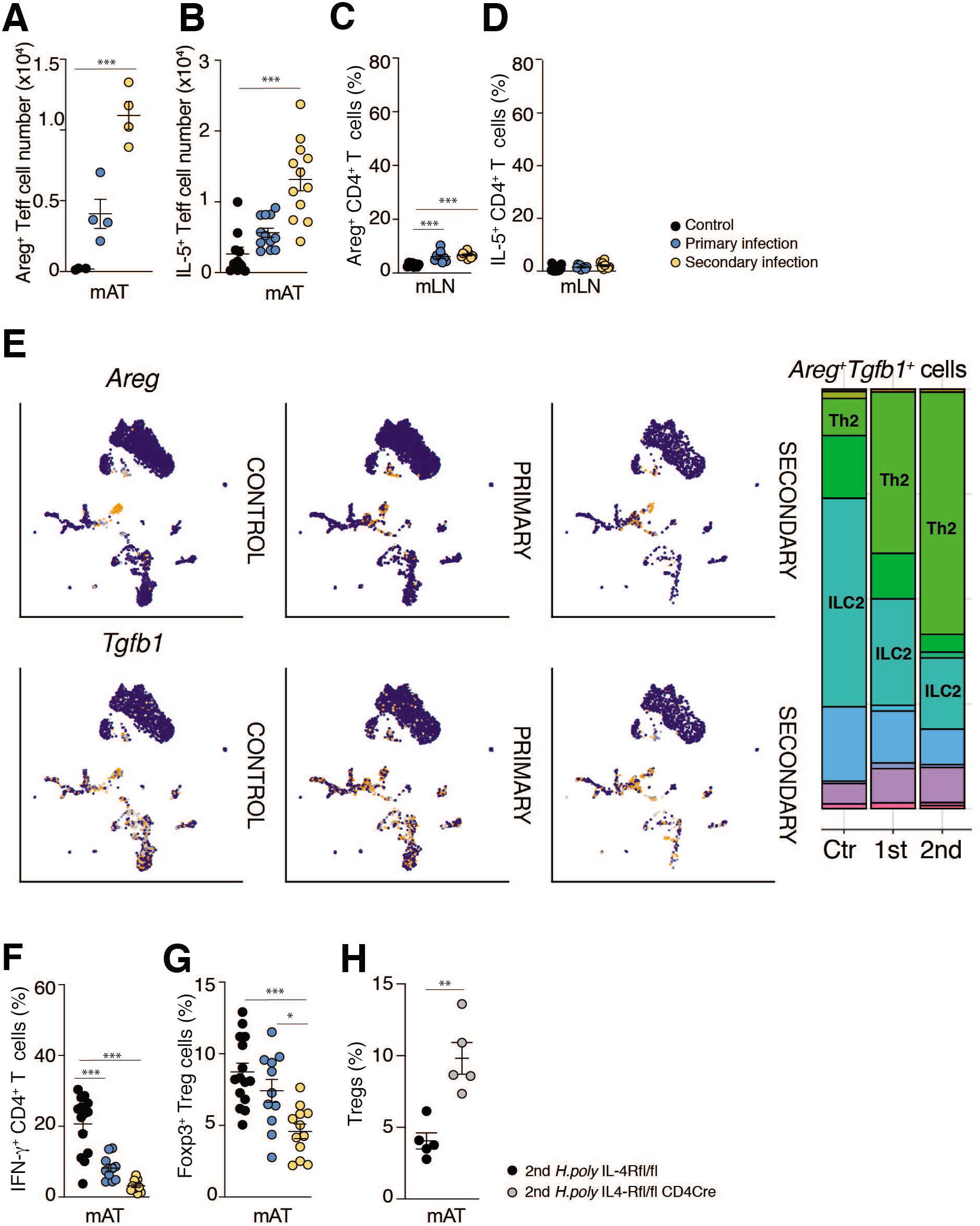
Th2_RM_ cell cytokine production in *H. polygyrus* infection. (**A, B**) Numbers of Areg^+^ (A) and IL-5^+^ (B) effector T cells in mAT in control and *H. polygyrus* infected mice. (**C, D**) Frequencies of Areg^+^ (C) and IL-5^+^ (D) effector T cells in mLN in control and *H. polygyrus* infected mice (gated on Foxp3^-^CD4^+^ TCRβ^+^ CD45^+^ live cells in lymphocyte gate). (**E**) UMAP of *Tgfb1* and *Areg* expression in cells from the scRNASeq SVF dataset as in Fig 2A split by experimental condition. Bar graph shows cluster distribution of double positive *Tgfb1* and *Areg* expressing cells (>0.5 normalized expression level) in scRNASeq SVF dataset. (**F**) Frequencies of IFN-γ^+^ effector T cells in mAT in *H. polygyrus* infection (as proportion of Foxp3^-^ CD4^+^ TCRβ^+^ CD45^+^ live cells in lymphocyte gate). (**G**) Frequencies of mAT Foxp3^+^ Treg cells as a proportion of CD4^+^ TCRβ^+^ CD45^+^ live cells in lymphocyte gate in control and *H. polygyrus* infected mice. (**H**) *Il4ra*^fl/fl^ *Cd4*-*Cre* and *Il4ra*^fl/fl^ mice were subjected to secondary *H. polygyrus* infection. Frequencies of mAT Treg cells (FOXP3^+^CD4^+^ TCRβ^+^ CD45^+^ cells in lymphocyte gate.) Dot represents BR and error bars represent SEM from BR (A-D, F-H). Data combined from 2-3 independent experiments (B-D, F, G), representative of 2 experiments (A, H), or from one experiment (E).

**Fig. S6.**
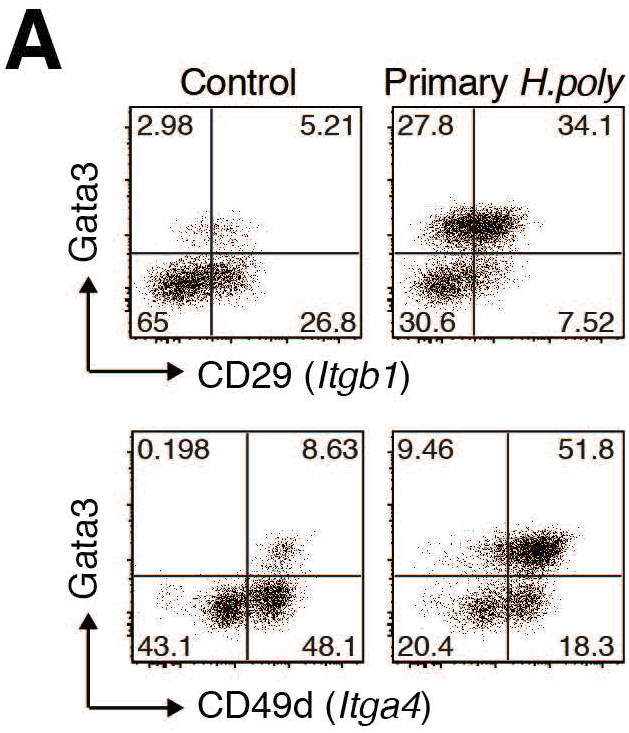
Integrin expression pattern on mAT Th2_RM_ cells in *H. polygyrus* infection. (**A**) Representative FACS plots of CD29 and CD49d expression by mAT Th2 cells in control and *H. polygyrus* infected mice (gated on Foxp3^-^ CD4^+^ TCRβ^+^ CD45^+^ live cells in lymphocyte gate). Data representative from two experiments.

**Fig. S7.**
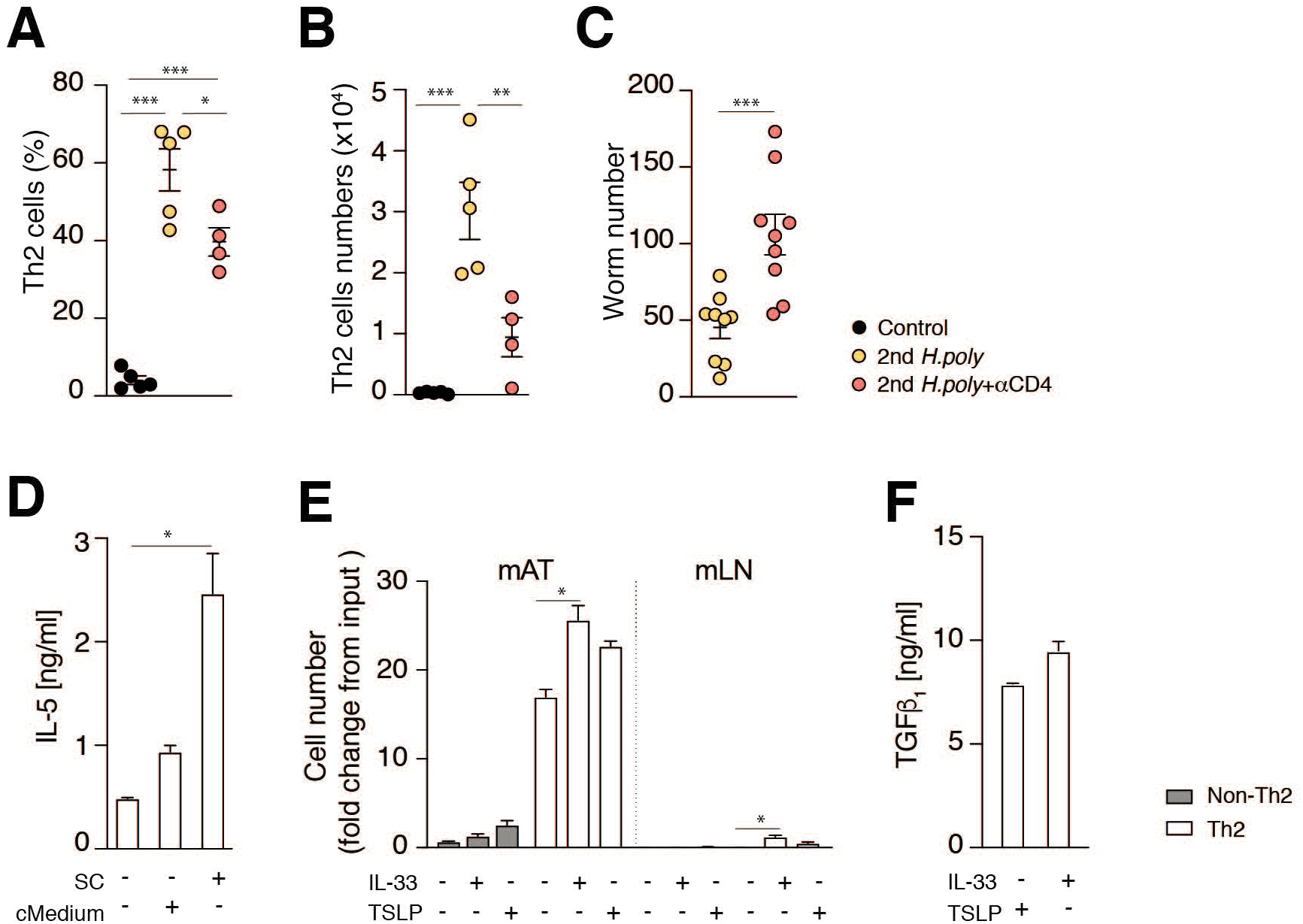
Interdependence of mAT Th2_RM_ and stromal cells. (**A-C**) Mice were subjected to secondary infection with *H. polygyrus* with experimental groups treated with anti-CD4 antibody as indicated (see Methods). Percentages (A) and numbers (B) of GATA3^+^ IL-33R^+^ Th2 cells (gated on Foxp3^-^ TCRβ^+^ CD45^+^ live cells in lymphocyte gate). (C) Worm counts from small intestine. (**D)** IL-5 levels from trans-well co-culture of sorted mAT Th2 cells (CD4^+^ TCRβ^+^ IL4-eGFP^+^ Foxp3RFP^-^) with SC isolated from *H. polygyrus* infected mice or supernatants collected from SC culture from *H. polygyrus* infected mice (cMedium), as indicated. (**E, F**) Th2 cells (CD4^+^ TCRβ^+^ IL4-eGFP^+^ Foxp3RFP^-^) and non-Th2 CD4^+^ T cells (CD4^+^ TCRβ^+^ IL4-eGFP^-^ Foxp3RFP^-^) were sorted from mAT and mLN of *H. polygyrus* infected mice and cultured for 3-6 days with addition of IL-33 [50ng/ml] or TSLP [50ng/ml] where indicated. (E) On day 3 cells were re-counted; numbers are presented as fold increases from the input cell numbers. (F) Levels of TGFβ_1_ in the supernatant from mAT Th2 cells after 6 days of culture. Dot represents BR and error bars represent SEM from BR (A-B) or from TR (D-F). Data combined from 2 independent experiments (C) or representative of 2 experiments (A, B, D, E), or from one experiment (F).

**Fig. S8.**
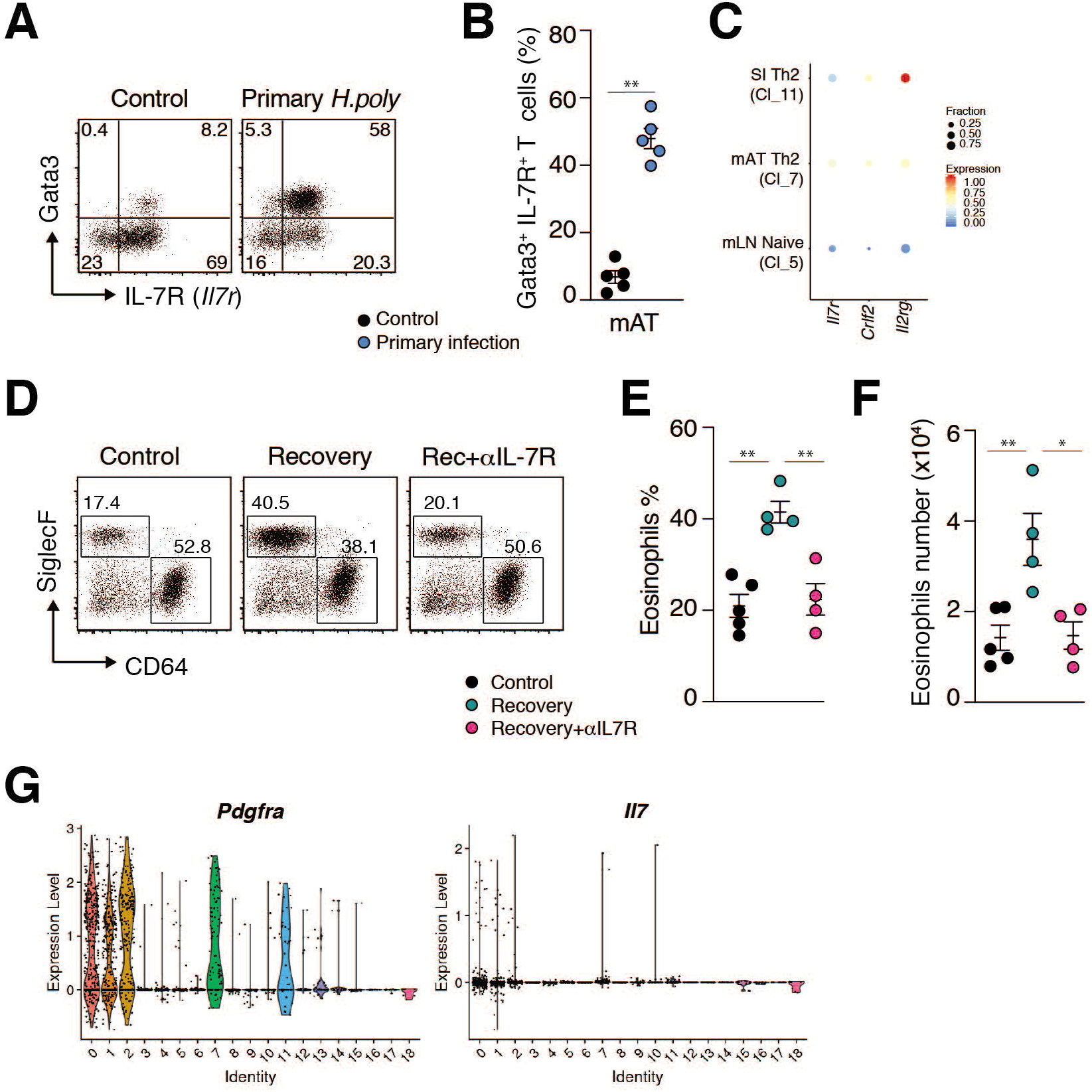
mAT stromal cells support Th2_RM_ population. (**A, B**) Representative FACS plots (A) and quantification (B) of in mAT GATA3^+^ IL7R^+^ Th2 cells (gated on Foxp3^-^CD4^+^ TCRβ^+^ CD45^+^ live cells in lymphocyte gate) during *H. polygyrus* infection. (**C**) scRNASeq dataset as shown in Fig.3: cluster-specific gene expression shown as dot plots where color represents the z-score of the mean expression across clusters and dot size represents the fraction of cells in the cluster expressing the selected gene. (**D-F**) TSLP signalling was blocked with anti-IL-7Ra antibody treatment during recovery after primary *H. polygyrus* infection (see Methods). Representative FACS plot (D) frequencies (E) and numbers (F) of SiglecF^+^ CD64^-^ eosinophils (gated on CD45^+^ CD11b^+^ cells) in mAT. (**G**) scRNASeq dataset with cluster identification as shown in Fig.2A: violin plot representing expression levels of *Pdgfra* and *Il7* according to cell cluster. Dot represents BR and error bars represent SEM from BR (B, E, F). Data from one experiment (A-G).

**Fig. S9.**
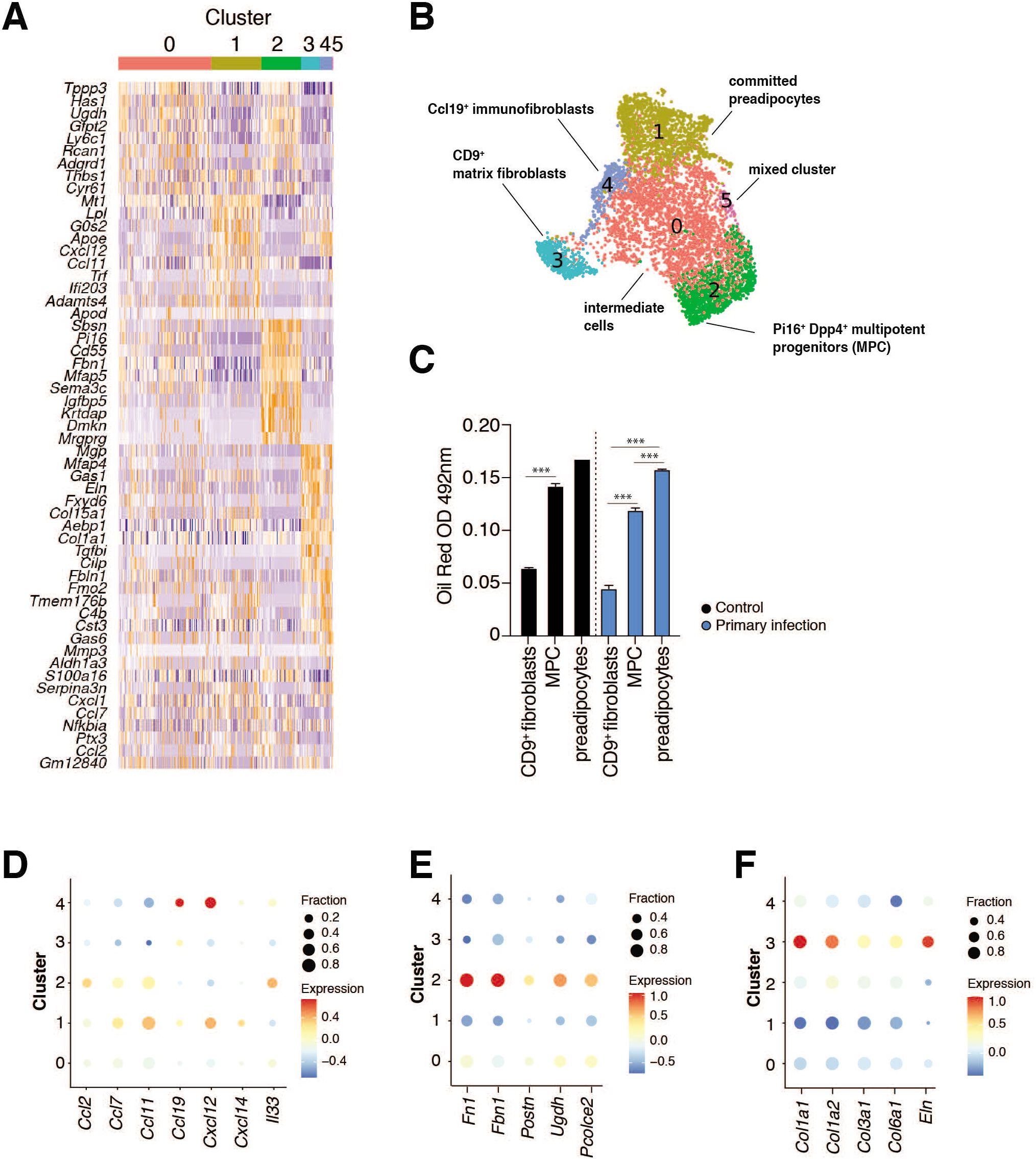
Characterization of mAT stromal cell populations in *H. polygyrus* infection. (**A**) Heatmap showing expression of top 10 differentially expressed genes in clusters as in panel B. (**B**) UMAP plot of 6059 mAT MSC from control (2457 cells), primary infected (2640 cells) and secondary infected mice (962 cells). Unsupervised clustering distinguished 6 cell clusters (C); plots are color-coded according to cell cluster. Identified cell population based on expression of following markers: committed preadipocytes (C1): *Icam1, Apoe, Lpl, Fabp4, Pparg*; pluripotent progenitors (C2): *Dpp4, Anxa3, Cd55, Pi16, Il33*; CD9^+^ profibrotic cells (3): *Cd9, Wnt6, Eln, Mgp, Col1a1, Col15a1*, immunofibroblasts (C4): *Cd9, Ccl19*. (**C**) mAT SC were sorted into Ly6c^-^ CD9^+^, Ly6c^+^ and Ly6c^-^ CD9^-^ subpopulations and subjected to adipogenic differentiation (see Methods). Lipid content was quantified with Oil Red staining. (**D-F**) Sample-specific gene expression shown as dot plots. Color represents the log2 normalized expression of the mean expression across samples and dot size represents the fraction of cells in the cluster expressing the selected gene. Error bars represent SEM from TR (C). Data from one experiment (A, B, D-F) or representative of two independent experiments (C).

**Fig. S10.**
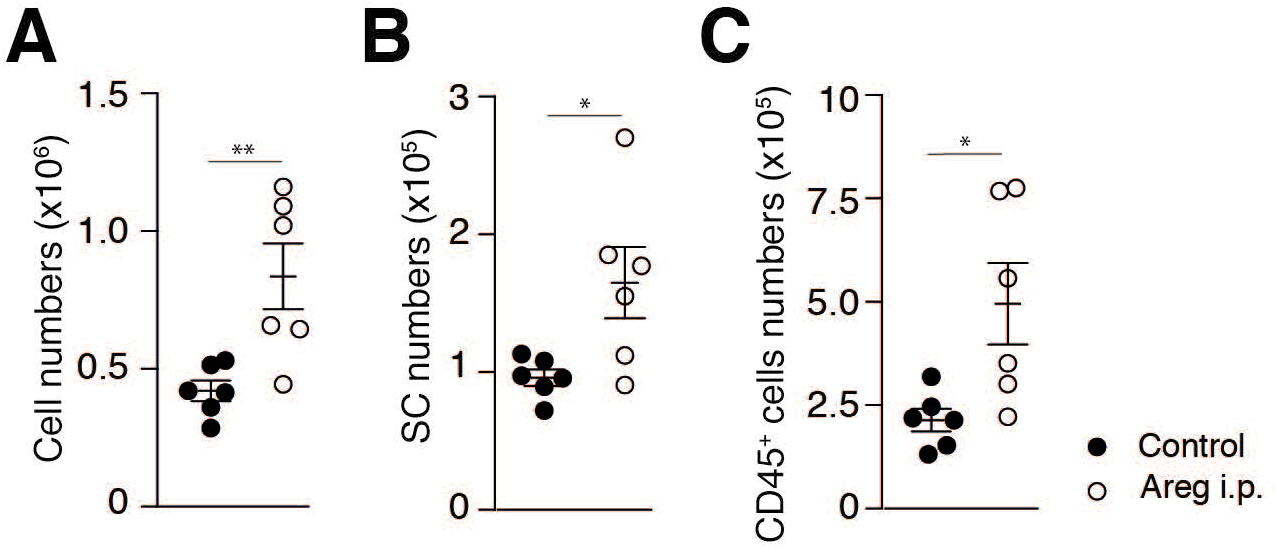
Areg supports mAT cellularity. (**A-C**) Mice were injected i.p. with Areg (3 injections, every third day) and mAT cells enumerated as indicated. Dot represents BR and error bars represent SEM from BR (A-C). Data from one experiment.

